# Therapeutic splice modulation of COL4A5 reinstates collagen IV assembly in an organoid model of X-linked Alport syndrome

**DOI:** 10.1101/2025.06.10.658776

**Authors:** Hassan Saei, Bruno Estebe, Nicolas Gaudin, Mahsa Esmailpour, Julie Haure, Olivier Gribouval, Christelle Arrondel, Vincent Moriniere, Pinyuan Tian, Rachel Lennon, Corinne Antignac, Geraldine Mollet, Guillaume Dorval

**Affiliations:** Laboratory of Hereditary Kidney Diseases, Inserm UMR 1163, Imagine Institute, Université Paris Cité, Paris, France; Necker Bioimage Analysis Core Facility of the Structure Fédérative de Recherche Necker, INSERM US24/CNRS UAR 3633, 75015 Paris, France; Department of Genomic Medicine for Rare Diseases, Necker-Enfants Malades Hospital, Assistance publique, Hôpitaux de Paris (AP-HP), Paris, France; Wellcome Trust Centre for Cell-Matrix Research, University of Manchester, Manchester, United Kingdom; Department of Paediatric Nephrology, Royal Manchester Children’s Hospital, Manchester University Hospitals NHS Foundation Trust, Manchester, United Kingdom

**Keywords:** Alport syndrome, mRNA splicing, ASO, Type IV collagen, *COL4A5*, Kidney organoid

## Abstract

Kidney organoids are an emerging tool for disease modeling, especially genetic diseases. Among them, X-linked Alport syndrome (XLAS) is a hematuric nephropathy affecting the glomerular basement membrane (GBM) secondary to pathogenic variations in the *COL4A5* gene encoding the α5 subunit of type IV collagen [α5(IV)]. In patients carrying pathogenic variations affecting splicing, the use of antisense oligonucleotides (ASOs) offers immense therapeutic hope. In this study, we develop a framework combining the use of patient-derived cells and kidney organoids to provide evidence of the therapeutic efficacy of ASOs in XLAS patients. Using multiomics analysis, we describe the development of GBM in wild-type and mutated human kidney organoids. We show that GBM maturation is a dynamic process, which requires long organoid culture. Then, using semi-automated quantification of α5(IV) at basement membranes in organoids carrying the splicing variants identified in patients, we demonstrate the efficacy of ASO treatment for α5(IV) restoration. These data contribute to our understanding of the development of GBM and pave the way for a therapeutic screening platform for patients.

## Introduction

Alport syndrome (AS) is the most frequent hereditary glomerulonephritis, due to a structural defect of the glomerular basement membrane (GBM)(1, 2). In the kidney, the main component of the basement membrane (BM) is type IV collagen which functions as a heterotrimer of α1α2α1(IV) predominantly in the early nephrons, α3α4α5(IV) mainly in the mature GBM but also in distal tubules and α5α5α6(IV) in the Bowman’s capsule and distal tubule BM(3),(2),(4),(5). The developmental switches in the GBM of laminin from α1β1γ1 to α5β2γ1 and of type IV collagen from α1α2α1(IV) to α3α4α5(IV) networks are crucial for maintaining the integrity of the glomerular filtration barrier(2, 6). The unique property of the collagen networks in BMs is the requirement for all specific chains of type IV collagen to be present in order to synthesize heterotrimers. If one chain is absent, the remaining intact chains are unable to form trimers and none of the trimer chains are expressed in the BM. The molecular network of these switches is not yet well understood(1).

AS is secondary to a defect in α3α4α5(IV) heterotrimers. Pathogenic variations leading to AS may involve the *COL4A3-4-5* genes, with autosomal recessive, dominant (*COL4A3-4*), or X-linked (*COL4A5*) inheritance(7, 8). A relative genotype-phenotype correlation has been described, ranging from long-life isolated hematuria to development of chronic kidney disease leading to kidney failure in the early adulthood in the most severe forms (mainly recessive or X-linked in males)(9–11). To date, the renin-angiotensin system blockers are the standard of care for all forms of AS, but they only delay kidney failure and no curative therapy exits to date.

We recently identified and validated splice-altering variations in patients with X-linked AS (XLAS) phenotype(12). In this study, we detected 14 pseudo-exon retention events in 19 patients with X-linked AS (XLAS). These variants are perfect targets of steric-block antisense oligonucleotides (ASOs), an emerging personalized therapy that can ideally mask the newly created splice sites or splicing modulator sites and restore wild-type RNA transcripts and normal protein expression. However, the lack of robust *in vitro* models is a limitation for preclinical evaluation of ASO capability to amend splicing.

Different studies have reported the generation of various XLAS mouse models, such as *col4a5^del-ATGG^*, *col4a5^tm1Yseq^* (p.G5X), and *col4a5^-/y^* (p.Arg471*) mice (13–15). These models can replicate phenotypic characteristics comparable to those observed in humans, such as GBM abnormalities, proteinuria, and renal failure. While using mouse models offers unique benefits, such as the ability to measure renal function, the validation of new therapeutic approaches poses challenges owing to the extensive time necessary for model generation and evaluation, as well as intrinsic differences in non-coding sequences between various strains of mice. As a result, it is necessary to develop more complex *in vitro* models that can rapidly mimic disease conditions and reflect the genetic background of the patients.

The emergence of induced pluripotent stem cell (iPSC) technology offers unprecedented insights into the previously inaccessible window for human organ development *ex utero*(16),(17),(18). The development of kidney organoids enables the study of kidney tissue organization and cell differentiation profiles over time(19, 20), as well as the development of kidney disease models (21–23). Although the differentiation of kidney organoids from iPSCs has evolved rapidly, it still requires optimization of the protocol to meet specific disease characteristics(17),(18),(24),(25). Organoid models of AS with missense or truncating coding variants have shown GBM defects(26, 27), offering the hope of better characterizing the pathophysiology leading to AS and testing therapeutic approaches.

In this study, we implemented a platform for the generation of iPSC-derived kidney organoids for the development of a preclinical framework encompassing the validation of efficient ASOs targeting various splicing events. By employing confocal imaging and multiomics validation, we embarked on the thorough characterization of iPSC-derived kidney organoids, describing crucial refinement in culture protocol for both cellular and extracellular matrix development. Optimization of the kidney organoid differentiation protocol and characterization of the organoids at different time points enabled us to better recapitulate and unravel XLAS disease with BM defects in kidney organoids harboring patient deep-intronic *COL4A5* variations. Overall, it produced a scalable model for ASO therapeutic approaches, allowing a two-step preclinical ASO assessment framework, utilizing first XLAS patient-derived primary cells and then progressing to kidney organoids featuring patient-specific variations, which opens the way for the development of precision therapy for patients with XLAS.

## Results

### Multiomics analysis of kidney organoids in prolonged culture revealed podocyte and extracellular matrix dynamics

Our initial goal before developing the disease model was to optimize culture conditions for accurately assessing the AS molecular phenotype. We maintained organoids in culture for 38 days and beyond to evaluate BM and podocyte molecular dynamics (classical culture terminates at day 22) (**Fig. 1a**). The integrity of distinct cell types within the organoids was assessed by immunostaining for podocyte marker WT1, proximal tubule epithelial cell marker LTL, and distal tubule epithelial cell marker CDH1 (**Fig. 1b**). Transmission electron microscopy (TEM) analysis of the organoids at day 38 of culture revealed podocyte assembly inside parietal-epithelial-like cells (PEC-like) and deposition of extracellular matrix components in the basal site as has been previously observed in organoids at day 22 of the culture(26, 28) (**Fig. 1c**). Multiomics (RNA-seq and proteomics) investigations on organoids at early (day 22), mid (day 32) and late (day 38 or 42) time points revealed podocyte and ECM associated gene and protein dynamics in culture (**Fig. S1a,b**). The complete list of the differentially expressed genes (DEGs) obtained from bulk RNA-seq analysis of organoids from day 32 vs day 22 and day 42 vs day 22 are provided in **Table S1**, and **S2** respectively. The list of proteins differentially regulated comparing late and early time points are provided in **Table S3** and **S4**.

**Fig. 1:**
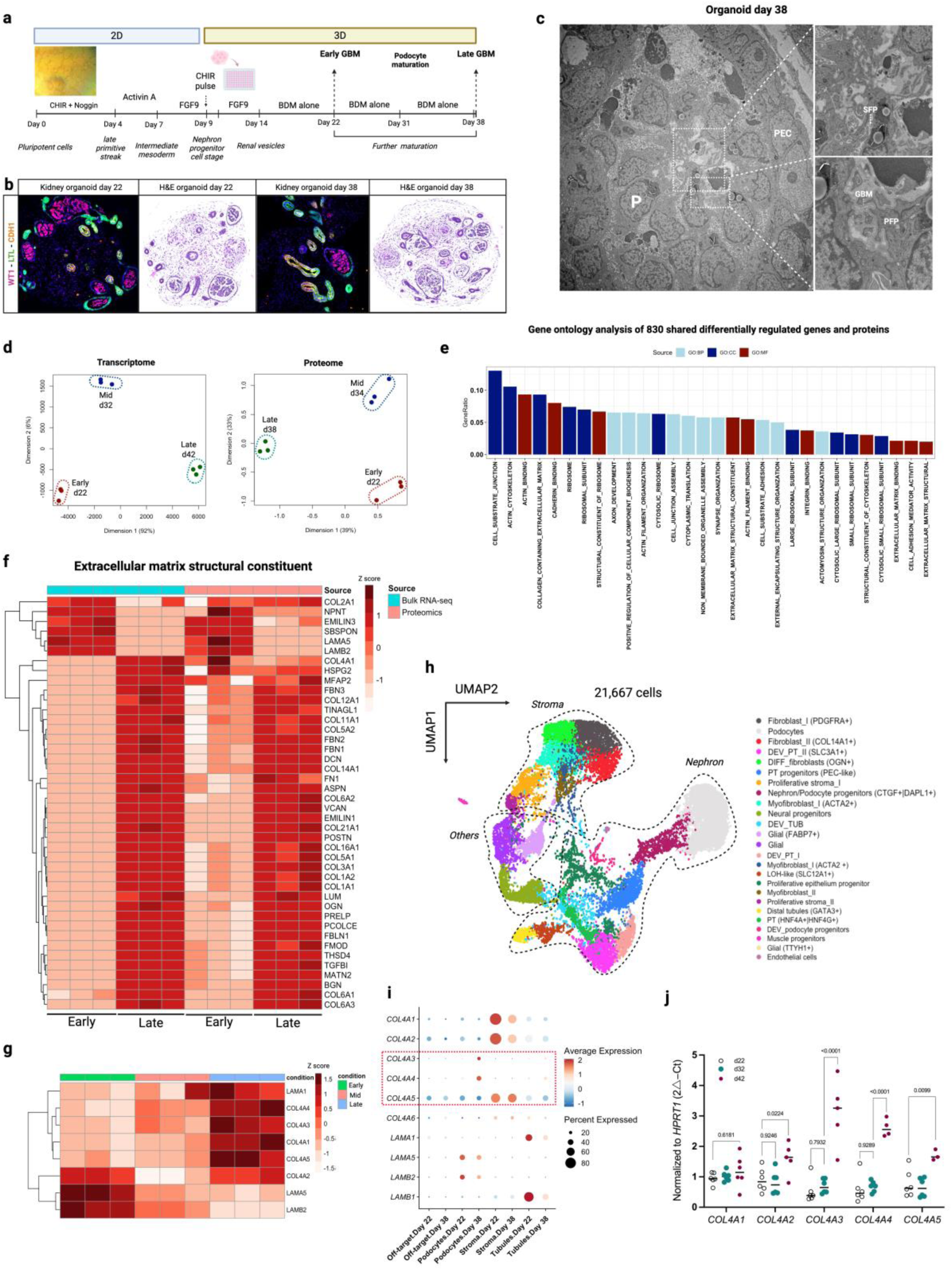
Multi-omics characterization of kidney organoids in prolonged culture. **(a)** Kidney organoid differentiation protocol transitioning from 2D to 3D cultures, with focus on prolonged culture. **(b)** Immunofluorescence and hematoxylin and eosin (H&E) staining of kidney organoids at day 22 and day 38, demonstrating their structural integrity. **(c)** TEM image of an organoid (day 38), highlighting glomerular structures with podocytes (P), primary and secondary foot processes (PFP, and SFP), glomerular basement membrane (GBM) and parietal epithelial cells (PEC) (magnifications: right image, 1250x; upper image, 4300x; lower image, 8500x) **(d)** MDS plot for transcriptomic and proteome data, showing distinct clustering of samples from different time points (early, mid, and late). **(e)** Gene ontology enrichment analysis (GOCC: cellular components; GOBP: biological processes; GOMF: molecular function) of 830 shared differentially regulated genes (fold change, 1.2) and proteins (FDR, 0.05) obtained from late versus early culture comparison. **(f)** Heatmap visualization of differentially expressed ECM structural genes and proteins overtime comparing early and late. Heatmap of type IV collagen and laminin genes expression from bulk RNA-seq are shown in panel **(g)**. **(h)** UMAP visualization of integrated single-cell RNA-seq data at day 22 and day 38, identifying distinct cell populations, including nephron population, stromal population and others (off-target population). **(i)** Dot plots of collagen type IV and laminin gene family expression across cell types and time points in the single-cell dataset. **(j)** RT-qPCR quantification of different chains of type IV collagen at kidney organoids harvested at early, mid and late culture confirming GBM maturation and dynamics in prolonged culture.

Multidimensional scaling plot (MDS) on bulk RNA-seq and proteomics data revealed a clear separation of samples from different time points and their replicates, indicating significant differences in gene and protein expression in organoids cultured for extended periods (**Fig. 1d**). We cross-matched transcriptome and proteome data (cellular fraction merged with ECM fraction) comparing late versus early time points and identified 830 shared differentially regulated biomolecules (genes and proteins) (**Fig. S1c,d**). Gene set over-representation ontology analysis (top 10 enrichments in each category, refer to **Fig. 1e**) revealed changes in genes and proteins associated with the extracellular matrix component and regulation (e.g., *FBN1, FBN2, VCAN, COL1A1, COL1A2, COL4A1, LAMB2*, *LAMA5*, **Fig. 1f**), cell junction assembly (e.g., *TJP1, CLDN3, CLDN11, ITGA2*, and *CDH11*, **Fig. S1e**) and podocyte development (e.g., *NPHS1, NPHS2, PODXL, ITGA3, LRP2*, and *PTPRO*, **Fig. S1f**). Notably, based on transcriptome data, the expression of *COL4A1*, *COL4A3*, *COL4A4, COL4A5,* and *LAMA1* was increased at day 42, while *LAMB2* and *LAMA5* expression was significantly decreased (**Fig. 1g**). Proteomics analysis was not successful in detecting collagen α3(IV) and α4(IV) specific peptides probably due to their minimal abundance compared to other type IV collagens, therefore it is not illustrated by the heatmap (**Fig. 1f**). Despite challenges in detecting collagen α3(IV) and α4(IV) peptides, these findings provide valuable insights into ECM composition dynamics over time, highlighting the unique molecular adaptations within long-term cultured organoids.

By performing single-cell RNA-seq on kidney organoids on days 22 and 38 (21,667 cells after integration), we aimed to verify which cell types were contributing more for the changes in ECM gene expression. We confirmed the existence of three main clusters within our organoids: nephron, stromal, and off-target clusters (**Fig. 1h**). Marker genes utilized to annotate each cell type are presented in **Fig. S2 and Table S5**. Focusing on podocyte populations, we observed different clusters including podocyte progenitors (expressing *DAPL1*, and *LYPD1*) and differentiated podocytes (expressing *DDN*). This indicated the presence of distinct podocyte populations within the organoids, each at different developmental stages. Gene expression analysis comparing two time points, with a focus on three main clusters, revealed an upregulation of GBM type IV collagen genes supporting GBM maturation and type IV collagen upregulation by day 38. Analysis of the podocyte cluster in our dataset revealed that *COL4A3* and *COL4A4* were overexpressed, whereas *COL4A1* and *COL4A2* mRNA expression decreased by day 38 in the podocytes (**Fig. 1i**). No significant difference in *COL4A5* expression was observed in podocytes when comparing day 22 to day 38. A slight decrease in *LAMB2* and *LAMA5* gene expression was also noted in podocytes at day 38, as shown by global transcriptome and proteome analyses. Type IV collagen expression was further studied by qPCR analysis of whole organoids harvested on days 22, 32, and 42, which confirmed the overexpression of the GBM type IV collagen network genes overtime (**Fig. 1j**).

We performed further organoid characterization through immunostaining for various proteins. Since we did not find the collagen α5(IV) antibodies to be specific, we inserted a V5 tag in exon 2 of the *COL4A5* gene, following the signal peptide. Therefore, labeling of the α5(IV) collagen in this study was always performed using anti-V5 tag antibody (**Fig. S3**). By co-labeling podocalyxin with type IV collagen, and tight junction protein 1 (ZO-1) with collagen α5(IV), we confirmed podocyte apical and basal polarity (**Fig. 2a,b**). In the kidney, α3(IV), α4(IV), and α5(IV) chains are produced by the podocyte, also by the distal tubular cells, and the α5(IV) chain is also produced by the PECs. Here, labeling of α3(IV), α4(IV), and α5(IV) collagens on days 22 and 38 revealed minimal deposition of the GBM collagen network, α3α4α5(IV), on day 22, with increased production and secretion of this network by day 38 (**Fig. 2c**). In addition, we also observed, as in the kidney, expression of the α5(IV) chain lining the cells surrounding the podocytes that we identified as PEC-like cells, since positively stained for claudin-1 (**Fig. S4a**), and of the α3(IV) and α4(IV) chains although almost undetectable at day 38. Further basement membrane analysis using laminin-specific antibodies (anti-Laminin-β1, Laminin-β2, and Laminin-α5) at different time points confirmed the establishment of a mature laminin network as early as day 22, which was expected since the laminin switch occurs earlier in the development than the type IV collagen switch(29) (**Fig. S4b**). Therefore, prolonged organoid culture is necessary for ECM maturation, particularly for the type IV collagen switch in the GBM, which is a key factor in the development of AS organoid models.

**Fig. 2:**
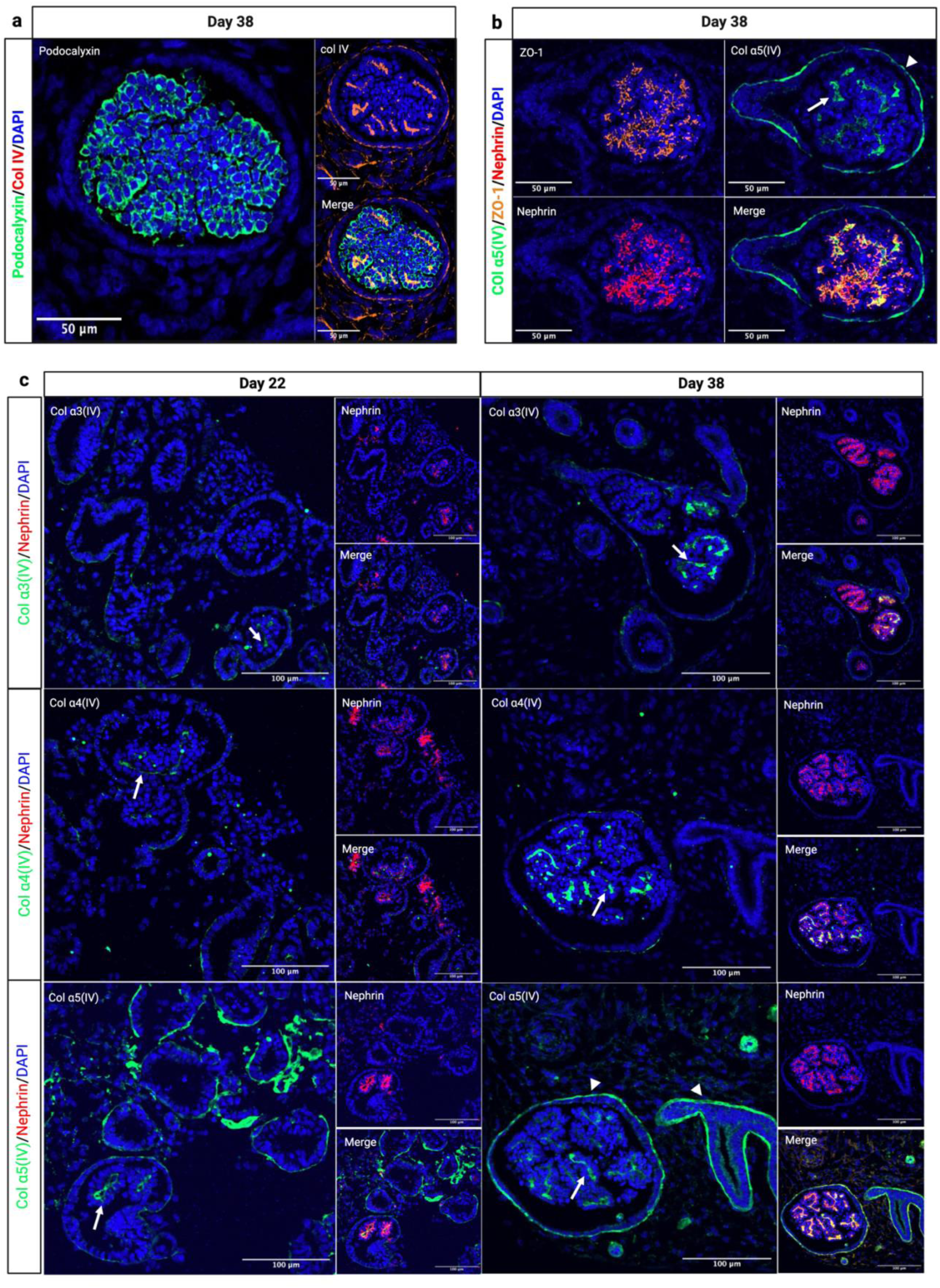
Immunofluorescence staining of kidney organoids highlighting the localization of various junctional and ECM associated proteins relevant to podocyte maturation. **(a)** Spatial distribution of podocalyxin (in green), and type IV collagen (in red) in organoid (day 38 of culture). Podocalyxin is mainly localized to the apical side of the podocytes, while type IV collagen (captures all chains α1-α6) is present in the basal side, forming the GBM. **(b)** Co-localization of tight junction protein (ZO-1) (in orange), nephrin (in red) and collagen a5(IV) (in green), within the basal side. **(c)** Type IV collagen network localization and abundance at days 22 and 38 of culture. This panel confirms the increased production of collagen α3(IV) and α4(IV) chains at day 38 and their deposition in the GBM (white arrows refer to α3α4α5 and arrow heads refer to α5α6α5 heterotrimers in the PEC-BM and TBM).

### XLAS organoid model with deep-intronic variations recapitulates basement membrane defects

For preclinical assessment of ASO therapy using the XLAS kidney organoid model, we generated two XLAS kidney organoid models by introducing deep-intronic variations in the *COL4A5* gene in the previously used control male iPSC line containing the V5 tag in the *COL5A5* gene. Both variations (c.277-560T>G identified in patient P3 and c.609+879A>G identified in patient P14) lead to creation of a new splice site and to intron retention (IR) but with a complete absence of the WT transcript for the former event and residual expression of the WT transcript for the later, which were thus expected to lead to a severe and a moderate model respectively (refer to KI paper for more information(12))(**Fig. 3a,b**). Targeted RNA-seq and fragment analysis confirmed aberrant *COL4A5* splicing in both XLAS organoid models. H&E staining revealed no phenotypic differences in differentiation between the models (**Fig. 3b,c**).

**Fig. 3:**
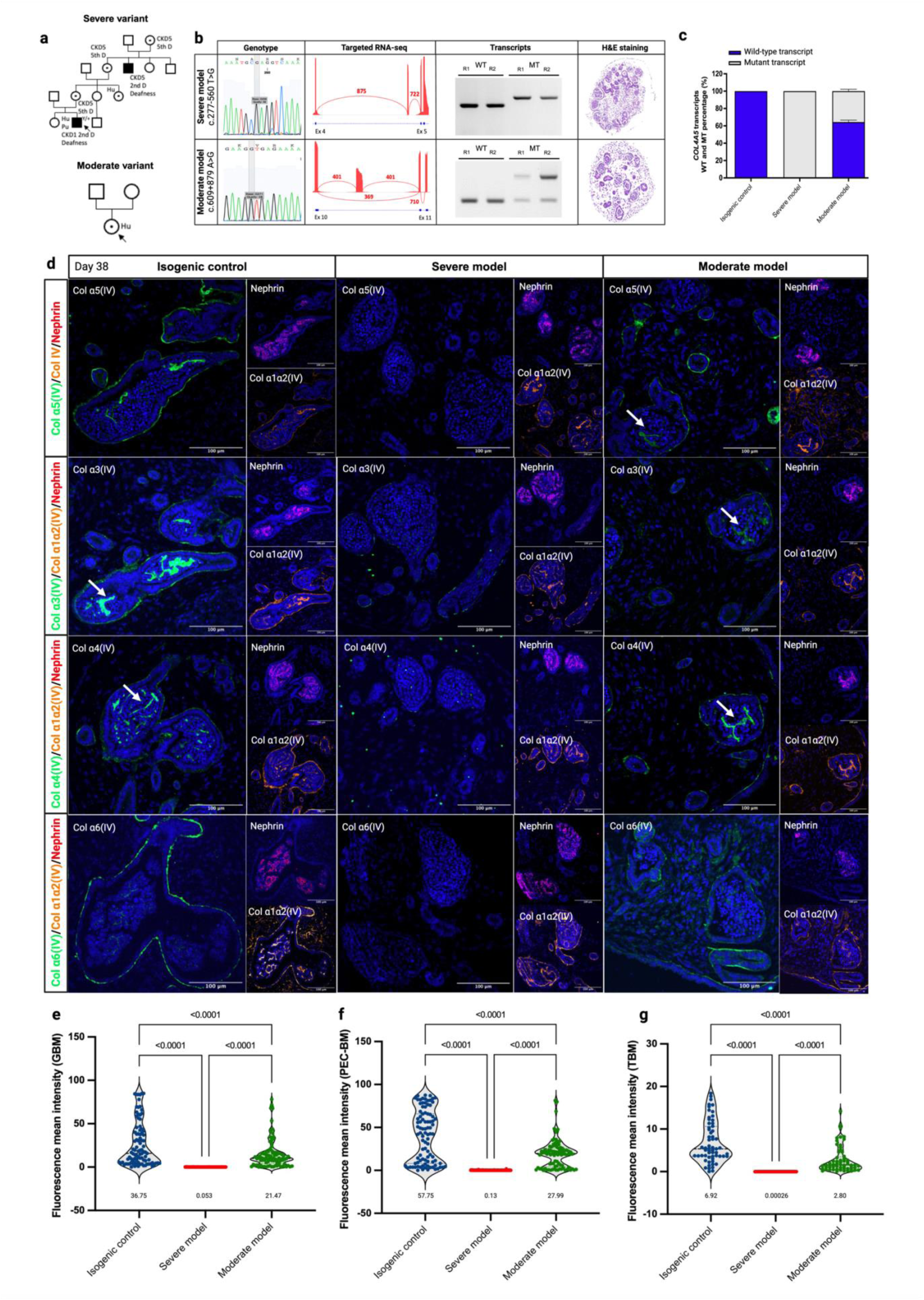
XLAS organoid model with deep-intronic variations recapitulating basement membrane disease. **(a)** Pedigrees and genotypes of the severe and moderate variants introduced in the control male iPSC line. **(b)** Multi level characterization of the molecular phenotype in organoid models using Sanger sequencing, targeted RNA-seq, RT-PCR results (R1, R2 indicates repeat 1 and 2) and H&E staining of XLAS organoids. Fragment analysis **(c)** along with targeted RNA-seq analysis of XLAS models revealed the absence of wild-type *COL4A5* transcript in the severe model and residual expression of normal transcript in the moderate model. **(d)** Immunofluorescence staining for collagen α3(IV), α4(IV), α5(IV) and α6(IV) chains in organoids (day 38), showed absence of type IV collagen networks (both α5α5α6 and α3α4α5) in the severe XLAS model and significant decrease in the moderate XLAS model. **(e), (f), and (g)** Quantification of collagen α5(IV) mean fluorescence intensity within GBM, PEC-BM, and TBMs in the severe, moderate and control organoids (n=60 object in each group and each object is either tubule or glomeruli) is shown (values below each condition indicate the mean intensity). No collagen α5(IV) secretion was observed in the XLAS severe models and significant reduction in collagen α5(IV) deposition was observed in all BMs in the moderate model. White arrows refer to the GBM.

Immunofluorescence staining of organoids at day 38 using type IV collagen chain-specific antibodies confirmed as expected the absence of collagen α3(IV), α4(IV), α5(IV), and α6(IV) proteins in the GBM, TBM and PEC-BM in the severe model and the qualitatively diminished presentation in the moderate model (**Fig. 3d**). To ensure reliable interpretation after therapy development, we developed a macro for FIJI (refer to methods) to semi-automate the quantification of BM proteins and perform a comparative analysis (**Fig. S5**). The mean fluorescence intensity of collagen α5(IV) in the GBM was 0.053 in the severe model, 21.47 in the moderate model, and 36.75 in the control (**Fig. 3e).** In the PEC-BM, the intensity was 0.13, 27.99, and 57.75 and for the TBM, the values were 0.00026, 2.80, and 6.92, respectively (**Fig. 3f,g**). This macro enabled accurate and rapid quantification of collagen α5(IV) in different BMs in the diseased and isogenic control organoids. Quantification of collagen α5(IV) mean fluorescence intensity revealed the absence of type IV collagen deposition in various BMs in the severe model. It also confirmed a significant decrease in signal intensity, indicating the defect in the moderate model (**Fig. 3e,f,g**).

Single-cell RNA-seq was performed on severe and moderate XLAS models on day 22 to determine the molecular aspects of the disease at early time point (**Fig. 4a**). Four single-cell experiment datasets were integrated (50,018 cells), and cell types were annotated using knowledge-based approaches and previously defined marker genes (**Fig. S2**). To identify differentially regulated genes in this experiment, we combined the two mutants into a single group, and performed differential expression testing using the DESeq2 method. This approach resulted in the identification of 391 DEGs (39 upregulated and 352 downregulated) (**Table S6**). As expected, the over-representation analysis of DEGs identified multiple pathways and ontology terms including, renal system/kidney development highlighting kidney-specific gene dysregulation. Moreover, the analysis revealed dysregulation of BM, extracellular matrix and cell-cell junction in diseased podocytes confirming the disease molecular pathology (**Fig. 4b**). Genes enriched in each ontology term are shown in **Fig. 4c**. Genes associated with the basement membrane and ECM, were either found to be overexpressed such as *ANXA1*, *S100A10*, and *HTRA1*, or downregulated (*HMCN1*, *COL23A1*, *LAMA1*, *ITGA8*, *COL13A1*, *SMOC1*, and *LAMA2)* in mutant podocytes (**Fig. 4d**). In the severe model we observed overall overexpression of *LAMB1* and *COL4A3* and *COL4A4* and in the moderate model we observed *LAMB2* and *LAMA5* overexpression. As expected, the *COL4A5* expression was downregulated in both XLAS models, due to the mRNA decay (**Fig. 4e**). Complete information of genes enriched in the other libraries are listed in **Table S7**. Overall, the single-cell RNA-seq experiment confirmed the enrichment of disease-associated pathways within the podocyte cluster, further supporting the relevance of organoids for XLAS modeling.

**Fig. 4.**
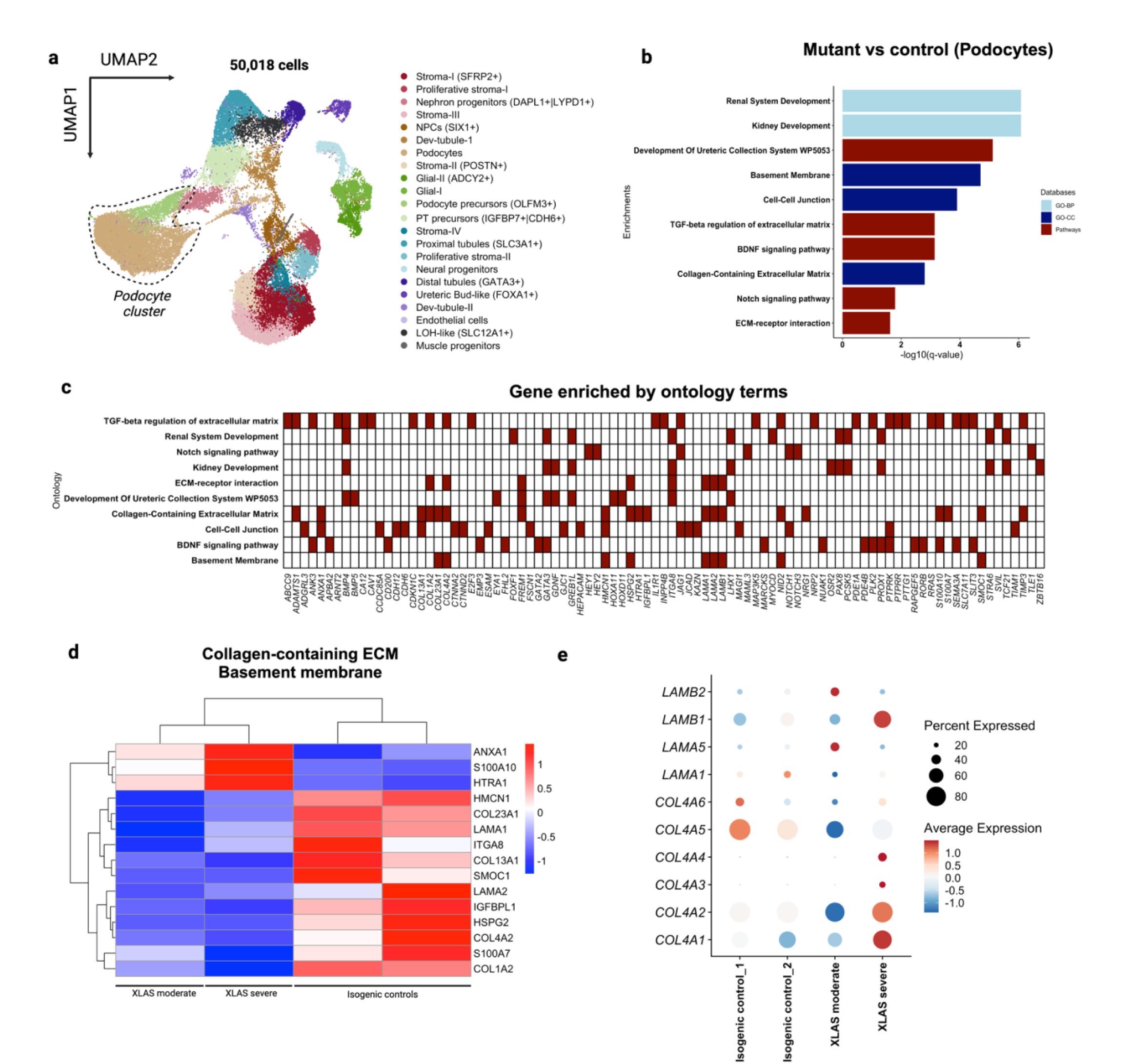
Single-cell transcriptional profiling in an XLAS organoid models revealed BM defects. **(a)** UMAP plot depicting 50,018 integrated cells across various cell types, including distinct podocyte clusters highlighted. **(b)** Gene ontology analysis comparing mutant (P3 and P14) versus control podocytes, showing enriched kidney and ECM-related pathways **(c)** Heatmap illustrating gene enrichment by ontology terms, with each row representing specific biological pathways (e.g., Notch signaling, kidney development), and each column denoting individual genes involved in these pathways. **(d)** Heatmap of BM and ECM-related genes in XLAS moderate and severe conditions versus isogenic controls, where red indicates higher expression and blue indicates lower expression. **(e)** Dot plot showing expression and percent expression of various type IV collagens and laminin genes across conditions, with dot size indicating percent expressed and color intensity representing average expression. The differences observed between isogenic controls 1 and 2 reflect transcriptional variations between differentiation batches.

### Validation of ASO amenability of splicing events in patient-derived primary cells and kidney organoids

As a first step we decided to validate the ASO-mediated restoration of *COL4A5* splicing in patient-derived fibroblasts. For this purpose, we selected six IR events observed in seven independent patients(12), P3, P10, P14, P15/P20, P16, and P19 (**Table 1**). Up to three ASOs were designed (refer to methods) to correct each IR event in mature *COL4A5* mRNA (the sequences and details of all ASOs are described in **Table 1**). We successfully corrected aberrant splicing in all (7/7) tested patient fibroblasts with deep-intronic variations in the *COL4A5* gene (for P3, and P14 refer to **Fig. 5a,f** and for P10, P15/P20, P16, and P19 refer to **Fig. S6**) employing lipofectamine-based ASO transfection. In particular, when testing three ASOs and their combination in fibroblasts of patients P3 and P14, we observed the best significant increase in the WT to mutant transcript ratio with ASO-M (0% in untreated cells to 73.9% in treated cells) and with ASO-A (69.26% in untreated cells to 96.46% in treated cells), in fibroblasts from patients P3 and P14 respectively (**Fig. 5c,h**), These results were confirmed by RT-qPCR (**Fig. 5e,i**) and targeted RNA-seq (**Fig. 5e,j**). This ASO screening is patient fibroblasts allowed us to select the best ASOs to be used in our severe and moderate XLAS kidney organoid models.

**Fig. 5:**
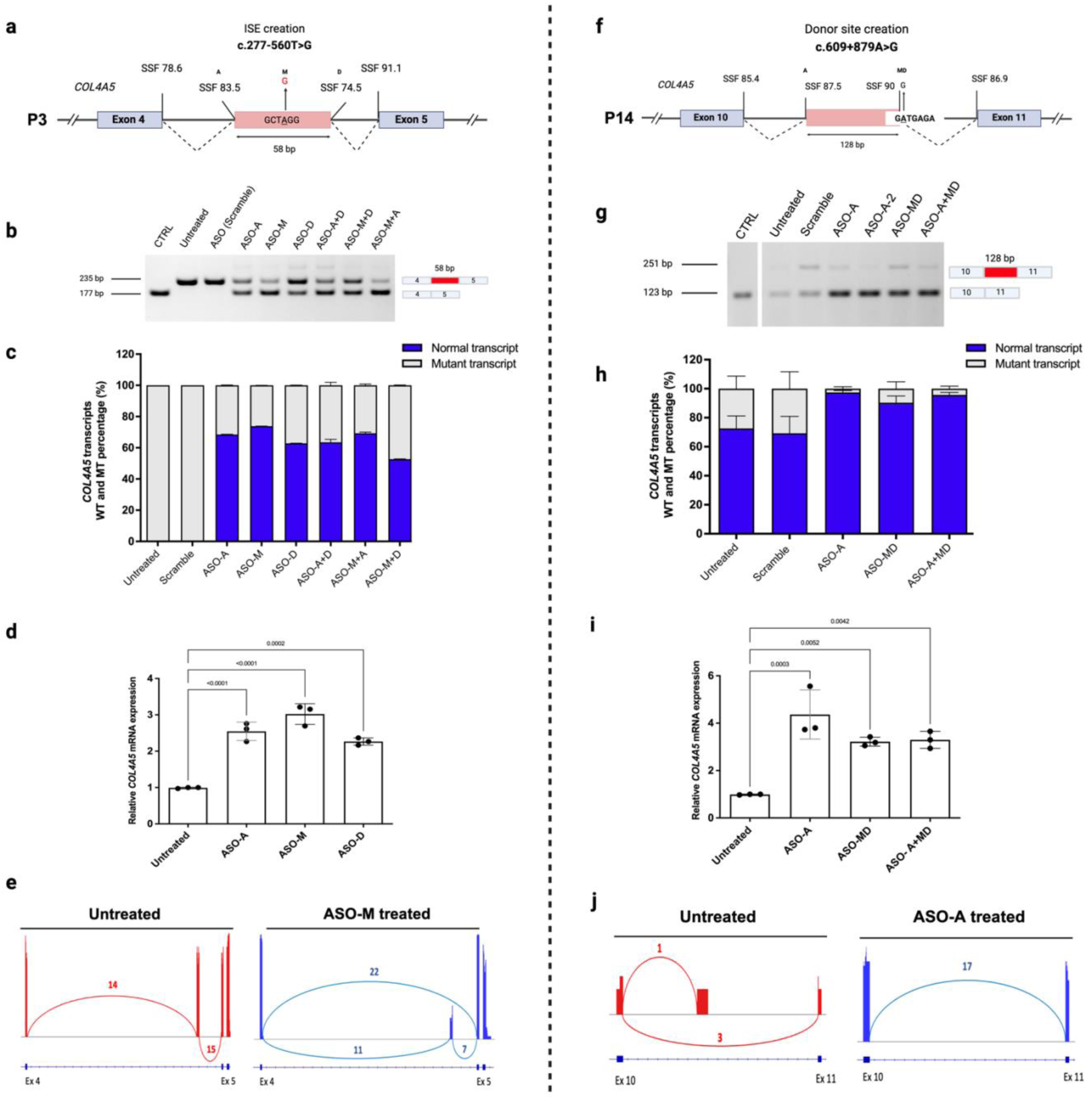
Evaluation of ASO efficacy in correcting splicing defects in patient-derived fibroblasts with deep-intronic *COL4A5* variants. **(a)** and **(f)** The schematic representation of the *COL4A5* variants identified inpatient P3 (c.277+560T>G) and P14 (c.609+879A>G), which create an intronic splice enhancer (ISE) and new donor splice site, leading to intron-retention. Splice Site Finder (SSF) score for each site is shown. **(b)** and **(g)** RT-PCR analysis of ASO treated and untreated cells, where ASO treatments reduced the levels of the mutant transcript and increased the proportion of the wild-type transcript. In P14, two faint bands are observed which are due to the NMD-mediated degradation of mutant mRNA and a low residual WT production. Panels **(c)** and **(h)** quantify the relative expression of wild-type and mutant transcripts across treatments by fragment analysis (n=3), showing the increase of wild-type to mutant transcript ratio post-ASO treatment. **(d)** and **(i)** RT-qPCR quantification of *COL4A5* mRNA normalized to *HPRT1*, show a significant increase after ASO treatments with Pvalue indicated above each bar plot (n=3). **(e)** and **(j)** The sashimi plots for untreated and ASO-treated samples, showing changes in splice junction usage before and after treatment.

**Table 1.**
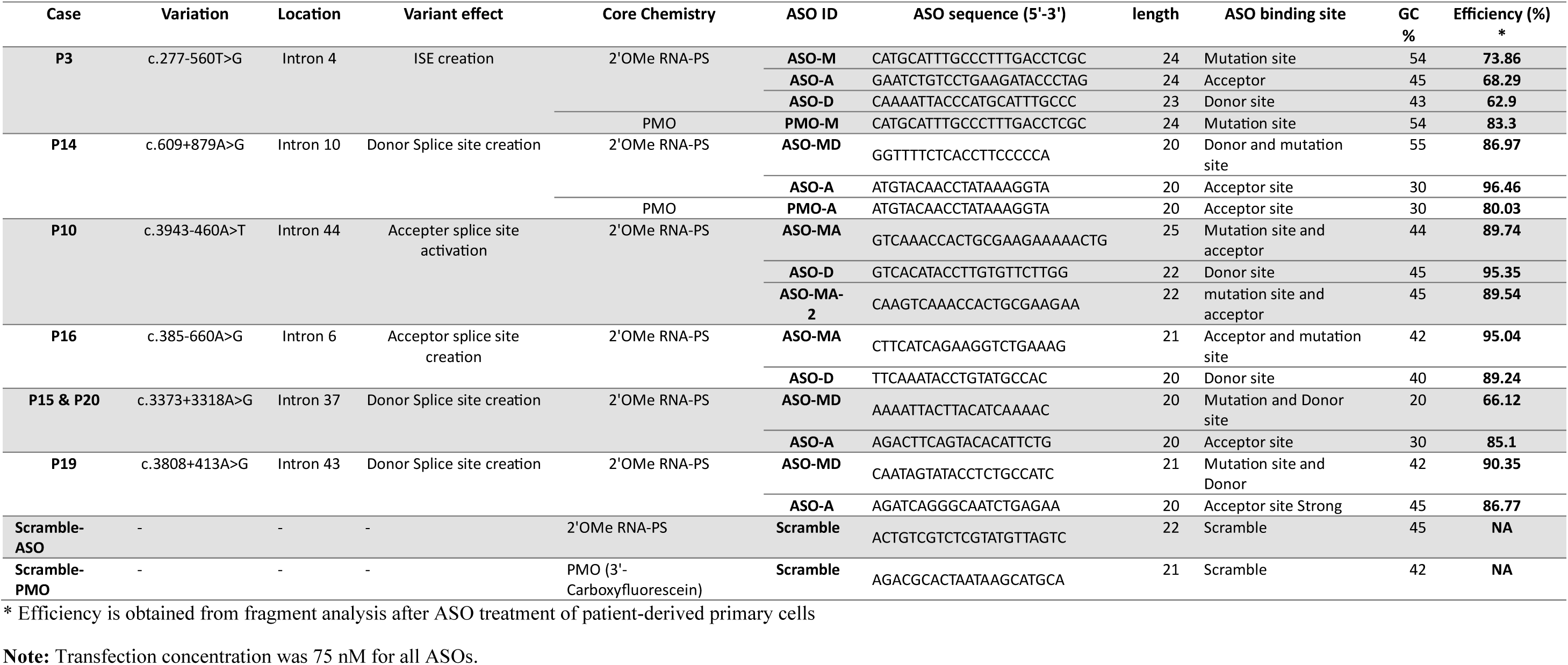
Variations and corresponding ASOs designed for splice modulation.

As our attempts of ASO transfection using Lipofectamine in kidney organoids was ineffective, we modified the oligonucleotide chemistry from 2’-OMe-PS RNA to phosphorodiamidate morpholino oligomers (PMO), which are neutral in charge and due to their chemical property can be more easily taken up by various cell types within the organoids. Kidney organoids were exposed to 5μM of PMO with the Endo-Porter system to increase endocytosis and PMO uptake. To assess the efficacy of PMO penetration within the organoid structure, fluorophore-labeled PMO was utilized (single transfection for three days at day 18). The untreated organoids exhibited no green fluorescence, confirming the absence of PMO (**Fig. 6a, top-right**). Conversely, labeled-PMO treatment resulted in positive signals from different cell types, mainly in PECs, tubular epithelial cells, and less in podocytes (**Fig. 6a, bottom panel**). Thus, we decided to use longer periods of treatment to improve the transfection in podocytes in the XLAS kidney organoids. The timeline is illustrated in **Fig. 6b**, with transfection starting on day 14 and renewed every 3 days until the final analysis on day 38. The experimental groups included isogenic controls treated with scramble-labeled PMO, severe XLAS, and moderate XLAS models and their respective PMO-treated counterparts.

**Fig. 6:**
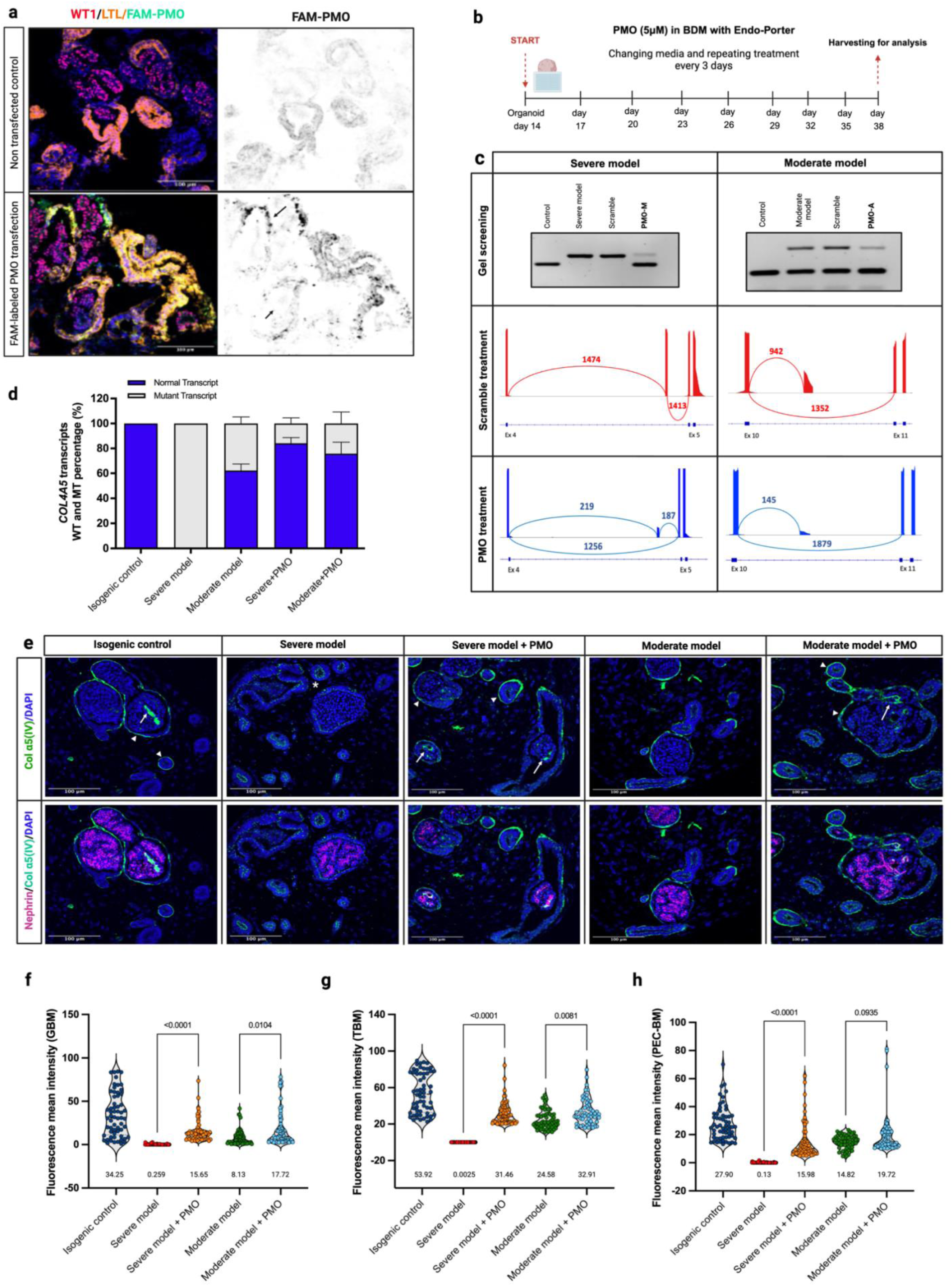
Restoration of basement membrane integrity in XLAS organoid models following antisense oligonucleotide therapy. **(a)** Immunofluorescence images showing PMO uptake (FAM-PMO, green) mainly in PEC-like cells and tubules LTL+ cells (yellow) after a single PMO transfection over three days (started at 10+8), demonstrating efficient uptake in tubular epithelial cells and PEC-like cells and less in podocytes (see black arrows). **(b)** The experimental timeline depicts the PMO treatment (5 μM) in organoids, starting at day 10+4, including media changes every three days and harvesting for analysis at day 38. **(c)** RT-PCR, and targeted RNA-seq results before and after PMO treatment of both severe and moderate models show a reduction of mutant *COL4A5* transcript and correction of splicing following PMO treatment in both models. **(d)** Quantification of normal to mutant *COL4A5* transcript ratio, shows a promising splice modulation capability of PMOs in organoids. **(e)** Immunofluorescence staining for collagen α5(IV), and nephrin, demonstrated significant restoration of collagen α5(IV) production and secretion as a network in the BMs of XLAS organoid models following PMO treatment (The white arrow refers to the GBM α5(IV) retention in the cells). In XLAS severe model before PMO treatment, α5(IV) protein was retained in the apical side of the epithelial cells. Restoration of collagen α5(IV) in the GBM confirmed the penetration of PMO in cells after three weeks of treatment. **(f), (g),** and **(h)** Quantification of collagen α5(IV) fluorescence intensity within the GBM, TBM, and PEC-BM showed significant increases in protein secretion and assembly following PMO treatment in both models (mainly severe model) compared to untreated controls. Values below each condition indicate the mean intensity. The * refers to the α5(IV) retention inside the cell.

The efficacy of PMO treatment was first evaluated by assessing effect on the expression levels of WT and mutant *COL4A5* transcripts in whole organoid lysates (**Fig. 6c**). In the severe model, the untreated samples displayed aberrant splicing, while PMO treatment led to a shift towards normal splicing, indicated by dramatically reduced aberrant transcript levels and the emergence of wild-type mRNA (0% WT junction before treatment – 85.15% after treatment). A splice modulation pattern was observed in the moderate model (58.93% WT junction before treatment – 92.83% after treatment), and PMO treatment significantly decreased the synthesis of mutant *COL4A5* transcripts (**Fig. 6c**, middle panel). Quantitative fragment analysis confirmed the significant increase in the WT to mutant ratio (80.33% in severe model, and 86.37% in moderate model) following PMO treatment (**Fig. 6d**). Immunofluorescence staining on organoids proved the competence of PMO treatment on the type IV collagen network deposition in different BMs in both severe and moderate XLAS models (**Fig. 6e**). In the untreated severe model, there was no α5(IV) deposition in BMs. PMO treatment significantly restored α5(IV) production, assembly with α3(IV) and α4(IV) and secretion into the GBM (mean intensity=15.65), TBM (mean intensity=31.46), and PEC-BM (mean intensity=15.98) in the severe model. Similarly, in the moderate model, untreated samples exhibited diminished α5(IV) expression (TBM=24.58, PEC-BM=14.82, and GBM=8.13), which was substantially improved post-PMO treatment (mean intensity in TBM=32.91, PEC-BM=19.72, and GBM=17.72). Despite observing PMO uptake mainly in PEC and tubular epithelial cells, the significant improvement in collagen IV deposition in the GBM of the severe model confirmed successful PMO uptake by podocytes. The quantitative analysis of mean fluorescence intensity (**Fig. 6f-h**), using our novel protein quantification method, showed significant increases in collagen α5(IV) deposition, particularly in the severe model and, to a lesser extent, in the moderate model.

Thus, the quantification and analysis of both collagen α5(IV) at the mRNA and protein levels before and after PMO treatment confirmed the effectiveness of using kidney organoids as scalable and robust *in vitro* models for developing ASO therapeutic approaches in XLAS.

## Discussion

Recent advances in integrative genome and transcriptome analyses have significantly improved the diagnosis of genetic diseases, particularly by identifying pathogenic variants affecting splicing. In total, it is estimated that around 20% of the variations identified in monogenic diseases are splicing-related variations(30). *In silico* scores have a low positive predictive value, and functional studies in valuable models are therefore essential. Moreover, steric-block oligonucleotide therapy has gained attention as an approach to treat various diseases caused by this class of variants(31). We and other groups have previously shown the importance of variants affecting *COL4A5* splicing in XLAS (intronic or exonic)(12)^-^(32–34). In this study, we developed a two-step preclinical ASO assessment framework, utilizing both patient-derived primary cells and kidney organoids featuring patient-specific variations. Then, using a multi-omics approach including single cell RNA sequencing and proteomics in combination with confocal imaging-based protein quantification, we have deeply characterized the development of the glomerular basement membrane of kidney organoids, wild-type or carrying the pathogenic *COL4A5* variants identified in two XLAS patients. We showed that prolonged culture of hiPSC-derived kidney organoids is crucial for the maturation of extracellular matrix components, especially in the GBM. Using ASOs, we then significantly re-expressed α5(IV) protein in the GBM of kidney organoids, thus opening the door to targeted therapy in these patients. The optimization of XLAS organoid development and splice modulation within 3D models emerged as a pivotal aspect of this study.

Organoids are an invaluable model for evaluating disease pathophysiology and therapeutic options, due to their genetic similarity to humans and extracellular matrix organization similar to kidney tissue(26),(27). Deep characterization of late matured kidney organoids following specific differentiations protocols is missing. Here, we employed multiomics (bulk and single-cell RNA-seq, and proteomics) and immunofluorescence staining on organoids harvested at early (day 22), mid (day 32), and late (day 38) time points. An integrative analysis of bulk RNA-seq and proteomics data confirmed dynamic changes in ECM and cell junction-related genes and proteins when comparing early (day 22) to late (day 38) time points(35). Notably, consistent with recent observations(36), *COL4A3*, *COL4A4,* and *COL4A5* expression was significantly enriched in organoids harvested at day 38 confirming the collagen switch during GBM development. Moreover, analysis of single-cell RNA-seq data (from day 22 and day 38) revealed upregulation of *COL4A3*, and *COL4A4* favoring GBM maturation. We identified distinct podocyte populations in the datasets, from early podocyte progenitors to “more” differentiated podocytes. By day 38, progenitor populations had decreased, giving rise to a more mature podocyte profile. Immunostaining analysis of type IV collagens in the GBM at early and late time points, confirmed increased mature collagen deposition, which was not investigated before. All these findings revealed the importance of maintaining organoids in culture for longer period to have more mature GBM, which is necessary for XLAS modeling.

Focused analysis of the podocyte cell population in the single-cell data identified pathways and ontology terms that could explain the cell-type specific pathology of the disease. This includes the evolution of the basement membrane, cell-cell junction and ECM receptor interaction in podocyte dysfunction. Proteins encoded by *ANXA1*, *S100A10*, and *HTRA1* genes play key role in cellular processes related to the ECM regulation and inflammatory responses. The upregulation of *ANXA1* and *HTRA1* could have a protective role in response to ECM stress and injury(37–39). On the contrary, overexpression of *S100A10*, which contributes to plasminogen activation and ECM remodeling, is more likely to promote ECM degradation(40). In disease podocytes, downregulation of structural integrity-associated genes affects both cell adhesion and ECM regulation, contributing to disease progression. *HMCN1*, encoding Hemicentin-1, is crucial for cell adhesion, and its reduced expression weakens podocyte junctions, leading to compromised cellular stability(41). Similarly, *ITGA8* downregulation loosens the attachment between podocytes and the basement membrane, aggravating podocyte injury and destabilizing the filtration barrier(42). Additionally, *SMOC1* downregulation limits its protective role in ECM organization and TGF-β signaling suppression, allowing profibrotic signaling to proceed unchecked(43). Together, the downregulation of these genes probably reflects both a loss of compensatory mechanisms, as seen with *SMOC1*, and contributions to disease pathology, as with *HMCN1* and *ITGA8* and contributes to the relevance of the model.

Patient-derived cell models most often allow analysis of transcription in the patient’s genetic background. This is only possible if the gene is expressed in accessible cells (which is the case for the *COL4A5* gene in fibroblasts, but not the case for *COL4A3-4* genes not expressed by fibroblasts, since only the α5α6α5(IV) chains are expressed at the dermo-hypodermal junction). Thus, the kidney organoid model adds an important dimension by measuring the impact on the protein encoded by this gene. In particular, in AS, the ability not only to study α5(IV) chain precise localization on a cellular level but to quantify its level, offers a perfect readout.

After demonstrating the capability of prolonged cultured kidney organoids to recapitulate key features of AS at the BM level, we sought to develop a framework for transfecting ASOs in kidney organoids in order to modulate *COL4A5* mRNA splicing. Different groups have proved the applicability of ASOs in organoid models. A recent study have used ASOs (1μM) for splice modulation in hiPSC-derived cortical and subpallial organoids and observed effects on calcium channel function and gene expression(44). Another study generated patient-derived organoid of pancreatic ductal adenocarcinoma and tested free uptake and transfection-based ASO treatment and concluded that transfection-based delivery is more effective than free uptake(45). In our study, transfection of ASOs without carrier (directly added to the media up to 10µM in concentration) and with carrier (either Lipofectamine 2000) was not efficient in penetrating the organoids. Therefore, we shifted towards the PMOs due to their neutral charge.

Recent researches on PMO-based splice modulation in organoid model offers valuable insight into their therapeutic potentials. Hall et al., generated colorectal cancer organoids from mouse models and patient samples and used PMOs to inhibit RNA splicing targeting KRAS isoforms (*KRAS4B*). This approach significantly decreased tumor growth by altering splicing in cancer stem cell pathways(46). Parfitt et al., used optic cup organoid model with *CEP290* variant and treated these organoids with PMOs targeting deep intronic variation(47). They treated their organoids every three days for 4 weeks beginning at week 13 of organoid development. This treatment led to significant increase in the WT *CEP290* transcript (70%). Other researchers generated retinal organoids from hiPSCs to induce exon skipping in the *PROM1* gene. PMOs were applied to these organoids at a concentration of 10 μM, with treatment conducted weekly over a four-week period, achieving around 57% exon skipping(48). These researches showed the possibility of using RNA- or morpholino-based oligonucleotides in organoid models. Consistent with these successful studies, our study demonstrated that PMOs are highly efficient in restoring correct *COL4A5* splicing in both severe and moderate XLAS organoids.

In 2020, Yamamura et al. demonstrated the applicability of ASOs to induce inframe skipping of an exon harboring a pathogenic stop variation in a mouse model of XLAS in mice(15). Despite successful restoration of collagen IV networks in the GBM and TBM, this treatment strategy aimed only to reduce a severe disease phenotype to a less severe one, since the full protein correction could not be achieved. However, splice-switching therapy offers the potential to completely reverse the phenotype by producing an intact, functional protein, when aimed at IR suppression. Our approach for this aim involved analyzing the splice site recognition sites and masking a few regions to find the best target with highest WT to mutant ratio after treatment while minimizing off-target. The effectiveness of our approach was proven through experiments conducted on primary cells then on organoids. Our results confirmed the efficacy of ASOs in restoring normal *COL4A5* splicing in multiple patient-derived models. Furthermore, our prolonged cultured kidney organoid model and the framework of analyses, including rescue by ASOs described herein, can ultimately be generalized to the α3(IV) and α4(IV) chains expressed by kidney organoids, but also to other key proteins of the podocyte or other kidney cells with specific membrane localization and in which we have identified intronic variations(49, 50).

While kidney organoids are invaluable models for studying disease mechanisms and therapeutic strategies, they still face some limitations. Firstly, a key limitation is the absent vascularization of kidney organoids, which remains a challenge in fully mimicking *in vivo* kidney properties. Vascularization is crucial for enhancing podocyte and GBM maturation, and can also improve the efficiency of ASO delivery to targeted podocytes. Encouragingly, recent advancements in organoid culture device such as flow systems, offer promising opportunities to overcome this limitation(51–53). Incorporating vascularization strategies could further refine our model, enabling a more accurate disease representation and therapeutic evaluation. Secondly, unlike animal models, kidney organoids cannot recapitulate proteinuria and are limited by certain physiological readouts. However, it has been proven that hematuria and proteinuria are not the only readout in patients and animal models of AS. Indeed, the correlation between the amount of collagen chain in the extracellular matrix and disease severity has been proven in AS, in particular in a recent paper in biopsies from patients with AS(54). The correlation between expression levels of α5(IV) and the associated clinical phenotype could in future make it possible to extrapolate the results obtained on organoids, particularly for assessing the efficacy of therapies, to the results expected in an animal model, or even in patients. This could avoid the need for systematic use of an animal model.

In this study we succeeded to generate a robust framework for modeling XLAS and evaluating ASO-based therapeutic interventions. Our findings highlight the utility of kidney organoids in bridging the gap between basic research and clinical application. As the field of organoid technology continues to advance, we anticipate that these models will play an increasingly pivotal role in the development of precision therapies for genetic disease.

## Methods

### Ethics statements and patient tissue collection

All patients gave written informed consent for collection of biomaterials. The research was performed in accordance with the Declaration of Helsinki on human experimentation of the World Medical Association, and it was conducted with the approval of the Comité de Protection des Personnes pour la recherche biomédicale Ile de France.

### Primary and hiPS cells culture and maintenance

Patient-derived fibroblasts were cultured at 37°C with 5% CO_2_ in RPMI 1640 (Thermo Fisher Scientific) with 10% FBS and 1% penicillin/streptomycin 100U/ml. In some cases, to inhibit nonsense-mediated mRNA decay, the cells were treated with Emetine 30μM for 16h and collected after washing with 1x PBS.

The hiPS cells were cultivated in mTeSR Plus (StemCell Technologies), on vitronectin XF (StemCell Technologies) or Matrigel hESC qualified (Corning) pre-coated plasticware. The passaging of the cells was made once a week manually under a stereomicroscope (cutting/scrapping) or with RelesR reagent (StemCell Technologies). We used Cryostor CS10 to freeze them (StemCell Technologies). The human iPS cell line IMAGINE004 was generated by the IPS facility of Imagine Institute from commercial male PBMC (Stemcell Technologies) using Sendai Virus procedure(55).

### CRISPR/Cas9 mediated knock-in in the *COL4A5* gene

A V5 tag was inserted at the beginning of exon 2 of *COL4A5* at position c.384 in the (IMAGINE004) iPS cell line and, subsequently patient mutations c.277-560T>G and c.609-879A>G were inserted into the resulting iPS cell line via CRISPR-Cas9 genome editing strategy. We chose the closest crRNAs to the variation site from the CRISPOR website (http://crispor.tefor.net/) and we designed 60nt ssODN with phosphorothioate modifications for each insertion. (**Tables S8**). In brief, 300pmol of gRNA (Alt-R CRISPR-Cas9 crRNA XT and Alt-R CRISPR-Cas9 tracrRNA ATTO 550) and 150pmol of Cas9 (Alt-R S.p. HiFi Cas9 Nuclease V3) were mixed to form ribonucleoprotein (RNP) complexes (IDT DNA technologies). Then, 600pmol of ssODN was added to the RNP mix. Prior to nucleofection, hiPS cells were pretreated with the apoptosis inhibitor Y-27632 (10μM) for 2h and further dissociated with Tryple Select (Thermo Fisher Scientific). Cells were nucleofected with the RNP complexes using the P3 Primary Cell 4D-Nucleofector X Kit-S and 4D-Nucleofector X-Unit (Lonza), using the program CA-137. The cells were then transferred to Matrigel-pre-coated 24-well plates and cultured in mTeSR-Plus medium with 10µM Y-27632. After 2 days, 10% of the transfected cells were sorted at 1 cell per well in CellAdhere Laminin-521 (StemCell Technologies) pre-coated 96 well plates. Cells were incubated in mTeSR Plus medium supplemented with CloneR2 (StemCell Technologies) for 3 days. We used Quick-extract DNA Extraction solution (Biosearch Technologies) and Sanger sequencing to identify clones of interest. The absence of genetic rearrangements subsequent to CRISPR-Cas9 genome editing technology was verified using an SNP array (Imagine Institute, iPS platform).

### Generation of kidney organoids from hiPS cells

We optimized a previously published protocol(18) to generate kidney organoids using hiPS cell lines. Human iPSCs (4,000 to 8,000 cells/cm2) were dissociated with Tryple Select, and plated onto Matrigel coated plates (Corning Matrigel Basement membrane matrix Growth Factor Reduced). We add Matrigel on the hiPSC one day after their seeding (0.125mg of protein/mL in mTeSR Plus) to induce the formation of iPSC spheroids during three days without any media change. Four days after seeding, the cells reached 40–60% confluency and differentiation was initiated (day 0) using Basal Differentiation Medium (BDM) (Advanced RPMI 1640, 1x L-GlutaMAX, and 0.5% penicillin/streptomycin (Thermo Fisher Scientific). Then, we used BDM supplemented with Noggin (PeproTech) and Chiron99021 (Axon Medchem) for four days, 4µM on the first day to reduce cell death then 8µM, then BDM with Activin A 10ng/mL (Peprotech) for the next three days. From days 7 to 14, FGF9 was added at 10ng/mL (R&D Systems), with media changes every two days. By day 9, we added 3µM Chiron99021 for 2h or 4h depending on the number of renal vesicles observed. For 3D culture, cells were dissociated by day 10 with Accutase (StemCell Technologies) and plated at 25,000 cells per well onto 96-well plates “ultra-low adherent” with U-shaped bottoms (Corning) in the BDM. Differentiation continued in the same way from day 10 to 42, with media change every two days.

### Histology and Immunostaining

Organoids were immersed in 4% paraformaldehyde (PFA) at 4°C for 20 minutes and washed twice with 1x PBS without calcium and magnesium, then stored in PBS at 4°C. To prepare formalin-fixed paraffin-embedded (FFPE) sections, the PFA-fixed organoids were embedded in 2% agarose gel, enabling them to be embedded in paraffin, and cut into 4-μm sections. For immunofluorescence microscopy, the FFPE tissue samples were subjected to a serial deparaffinization process, involving two 10-minute deparaffinization steps, two 5-minute in 100% ethanol, 5 minute-in 90% ethanol, and an incubation in 80% ethanol, followed by washing in Milli-Q water. FFPE samples underwent heat-induced antigen retrieval by immersion in 10mM sodium citrate buffer (pH 6.0) in a rice cooker for 20 min. Then, they were treated with a 0.1M glycine/6M urea solution for 30 minutes at room temperature. The deparaffinized sections were subjected to nonspecific binding site blocking with 1x Casein in PBS and treated with primary and secondary antibody solutions (**Table S9**). The slides were mounted using Mowiol and analyzed using a ZEISS Z1 Spinning disk confocal microscope at the Imagine Institute Imaging platform. For protein quantification in organoids, glomeruli and tubule annotations were made and classified using QuPath (v0.5.1)(56) and sam-api (v0.6.1)(57) and manually corrected. These annotations were then exported a regions of interest (ROI). Then α5(IV) positive signal in each image was identified using Shallow Learning pixel classification with Ilastik (v1.4.0)(58). We finally measured α5(IV) mean intensity in glomeruli border and interior, using the ROIs and the Ilastik pixel classification models generated before, with a FIJI (v2.14.0)(59) macro. Refer to the GitHub page for more information (https://github.com/hassansaei/BMQuant).

### RNA extraction, cDNA synthesis and RT-qPCR

Total RNA was extracted from both organoids and primary cells using the Rneasy Mini Kit followed by DNase treatment. Single-strand complementary DNA (cDNA) was synthesized from 500ng or 1000ng total RNAs using SuperScript-II Reverse Transcriptase (Thermo Fisher Scientific). RT-qPCR analysis of target genes was performed using iTaq Universal SYBR Green Supermix (Bio-Rad) on a Bio-Rad CFX384 instrument. The relative gene expression was calculated using 2^-ΔΔ**CT**^ method. The primers used for RT-PCR and RT-qPCR analysis were provided in **Table S10**.

### Transcript analysis using FAM-labeled primers

For conventional PCRs involving gel screening and fragment analysis, a pair of variant-specific primers with forward primers labeled with 6-FAM were utilized. The optimal number of 30 PCR cycles for patient fibroblasts and organoids was established using qPCR. Fragment analysis was performed with 1μl of PCR product using ABI 3500xL fragment analyzer (Applied Biosystems). The area under the curve value was calculated using Peak Scanner software (Thermo Fisher Scientific) to determine the quantity of *COL4A5* WT and mutant transcripts.

### Targeted RNA sequencing

To prepare libraries for targeted RNA-seq, first-strand cDNAs was synthesized from total RNA using the SuperScript VILO kit (Thermo Fisher Scientific), and the second-strand cDNA synthesis kit (Thermo Fisher Scientific) was used to produce the second strand. The double-stranded cDNAs were purified using AMPure XP Reagent (Beckman Coulter), and 50ng of cDNA was used to prepare the NGS targeted RNA-seq libraries using a TWIST technology. The capturing probes were synthesized from a long-range PCR on the *COL4A5* full coding sequence in the plasmid pNLF1-C_hCOL4A5-Nluc (Addgene)(60). Fragmentation of cDNA was carried out using the Twist Library Preparation EF kit and Twist Universal Adapter System kit and biotinylation of the probes was performed using an in-house protocol. Paired-end sequencing (150bp+150bp) was performed using a MiSeq sequencer (Illumina). The FASTQ files were aligned to the GRCh37 assembly of the human genome using Hisat2 software, and the visualization of the junctional reads was performed using the sashimi plot view feature of the IGV software (Broad Institute).

### Bulk RNA sequencing and data analysis

Total RNA was extracted from the pool of nine organoids per condition (three time points: days 22, 32, and 42 with three replicates in each). The quantity and quality of the extracted RNAs were assessed using the Lunatic nucleic acid quantification system (Unchained Labs) and an Agilent Technologies 5200 Fragment Analyzer. The library was constructed with ∼200ng of total RNA using the Universal Plus mRNA-Seq Kit (Tecan). The 9 libraries were sequenced on the NovaSeq 6000 platform (Illumina) using the S1 flow cell at the genomics platform at the Imagine Institute. The fastq files were aligned to the Ensembl GRCh38 assembly of the human reference genome using the Illumina DRAGEN Bio-IT Platform (v4.0) and the gene count matrix was generated using the FeatureCount software(61) in R. The matrix of counts was analyzed using three independent statistical methods: edgeR(62) (v3.26.8), Deseq2(63) (v1.24), and limma-Voom(64) (v3.40.6). The relevant genes were identified from each comparison group using an absolute logarithm fold change (logFC) greater than 1.2 and the adjusted p-value (p<0.05). Gene enrichment or over-representation analysis was conducted in R using the enrichment function of the ClusterProfiler(65) package. Heatmaps and regular and bar plots were generated in R using pheatmap, tidyverse and ggplot2 packages, respectively.

### Sample enrichment for proteomic profiling

Kidney organoids harvested at days 22, 32, and 38 of differentiation (n=16 organoids pooled) subjected to cellular soluble protein and matrix protein enrichment, as previously described(3). Samples were washed three times with 1x PBS, then homogenized in Tris-lysis buffer (10mM Tris pH 8, 150mM sodium chloride, 1% Triton X-100, 25mM EDTA in LC-MS grade water, and an EDTA-free Roche complete protease inhibitor cocktail). The organoids were incubated in this solution for 1h at 4°C with constant rotation and centrifuged at 14,000g for 10 min at 4°C. The supernatant was then collected and labeled as fraction 1 (enriched with soluble cellular proteins). The pellet from the first step was resuspended in an alkaline detergent buffer (0.5% Triton X-100, 0.1M PBS, and 20mM ammonium hydroxide). The fraction 2 which contained the cell surface and transmembrane proteins was collected following the same process as fraction 1 (incubation and centrifugation to obtain the supernatant). The remaining pellet was resuspended in 50 µL of TEAB/SDS lysis buffer (100mM TEAB pH 8.5, 10% SDS), and the samples were subjected to sonication using Covaris AFA glass tubes in the Covaris LE220+ system to obtain fraction 3 (ECM fraction). The sonication parameters for kidney organoids were set as follows: 40s per sample, peak power of 500 watts, duty factor of 20%, 50 cycles per burst, and average power of 100 watts. Fractions 1 and 3 were analyzed separately. To reduce the samples, 5μl of a 100mM dithiothreitol solution was added to 100μl of the soluble fraction, and heated 10 minutes at 60°C. After cooling, 15μl of 100μl iodoacetamide solution was added to the fractions for 30 min at 4°C to alkylate the proteins. Sample preparation for label-free mass spectrometry-based proteomics involves overnight digesting samples with Trypsin Gold (Promega) in S-Trap Spin Columns, followed by purification using OLIGO R3 Reverse beads in acetonitrile. The resulting peptides were dried and subjected to mass spectrometry data acquisition using a Thermo Scientific Q Exactive HF hybrid quadrupole-Orbitrap mass spectrometer with a Dionex Corporation UltiMate 3000 Rapid Separation LC nanosystem at Manchester University.

### Bioinformatic analysis of the proteomics data

We utilized MaxQuant (MQ) software (v2.4.9) on the biocluster server at the Imagine Institute, which was operating under dotnet version 3.0, to analyze the raw proteomics data. Separate MQ parameter files (mqpar.xml) were generated for the cellular fraction and ECM data, with carbamidomethylation of cysteine as a fixed modification, oxidation of methionine, proline, and lysine, and N-terminal acetylation as dynamic modifications. These modifications were added to improve the collagen peptide identification. The MS data were searched against UniProt and TREMBL databases for humans (txid, 9606). MQ produces a separate peptide file for each analysis, including cellular fraction and ECM fraction. Perseus software was used to merge cellular with ECM peptides and data normalization and statistical analysis was performed. A MDS plot was generated to perform a quality check. Differentially expressed proteins were filtered based on the adjusted p-value (adjPval<0.05) and the differentially regulated proteins were visualized in R using ggplot2 function.

### Organoid dissociation and single-cell RNA sequencing

Two separate batches of sample collection were performed in this project. In the first batch, 16 organoids were collected on day 22 of differentiation for isogenic control organoids, severe XLAS organoids, and moderate XLAS organoids, resulting in three libraries. An additional isogenic control was harvested on day 22 and another one from the differentiation batch was kept in culture for 38 days and then harvested at this later time point (two additional libraries, five libraries in total). After a quick wash in PBS 1X, harvested organoids were dissociated by several rounds of 10 minute incubation at 37°C with 500µL of TrypLE, with homogenization by gently flicking the tube every five minutes. At the end of each incubation, the supernatant was aspirated, inactivated in PBS containing 10% SVF on ice and replaced by fresh TrypLE.

Mechanical dissociation was applied by pipetting up and down 5 times from the third incubation round. Finally, all collected supernatants were put through a 40µm cell strainer, and the dissociated cells were pelleted at 300g for 3 min, suspended in 1mL of PBS (1% BSA, 1mM EDTA) and placed on ice. Cell counting evaluated cell viability between 70% and 85%. Libraries were generated using Chromium Next GEM Single Cell 3’ and Next GEM Chip G Single Cell Kits (10x Genomics) at the genomics platform at the Institute du Cerveau. The quality of the libraries was assessed using the High Sensitivity D5000 ScreenTape Assay for TapeStation Systems (Agilent), and the libraries were sequenced on NovaSeq X using the NovaSeq X plus 10B Reagent Kit (100 cycles).

### Single-cell RNA-seq data analysis

The sequencing data were preprocessed using the CellRanger pipeline (10x Genomics, Cellranger count v7) with default parameters, aligned to GRCh38, and the resulting matrix files were analyzed using the Seurat (v5.1.0) package in R (v4.3.3) software. Outliers were detected and filtered using a function that defined the Median Absolute Deviation (MAD) with the number of mad (nmad) set to 5 for nCount_RNA and nFeature_RNA (both log1P) and 3 for the percentage of mitochondria. The SoupX(66) algorithm was used to remove ambient RNAs, and the DoubletFinder(67) algorithm was used to remove potential doublets. The Seurat object’s assay version is set to "v3" to utilize older reduction and data integration methods (CCA and RPCA). Integration features (nfeatures=3500) were selected, followed by the application of PrepSCTIntegration. Next, the integration anchors were identified using both CCA and RPCA reduction methods, and the datasets were integrated using the IntegrateData function. Dimension reduction was performed using RunPCA, and clusters were determined using resolutions, from 0.5 1.6. RunUMAP functions were applied to the integrated datasets, and the best resolution was selected based on the number of unique informative clusters. To compare organoids at two time points (d22 and d38), we applied the CCA method and uMAP with a resolution of 1.2. The scCustomize library was used to select a broader range of palettes for data visualization. The final cell-type annotations were added using knowledge-based methods, which involved extracting differentially regulated markers (using FindMarker) for each cluster and incorporating literature-based annotations(68, 69). To compare mutants with the isogenic control, we used CCA integration with tSNE visualization of clusters identified with a resolution of 1.6. The DESeq2(63) method was applied to a data frame obtained by subsetting the podocyte clusters and pooling the reads in each sample using the AggregateExpression function. We used the DotPlot function in the Seurat package to produce dot plots. Heatmaps were generated using the Pheatmap function.

### Designing Antisense oligonucleotides for splicing modulation

Antisense oligonucleotides (ASOs) with 2’-O-methylation (2’-Ome)-RNA chemistry and a full phosphorothioate (PS) backbone were designed and synthetized by Eurogentec using reversed-phase HPLC purification. The ASO binding location was chosen and the final sequence was determined by the probable off-target score obtained from the NCBI blast engine. The ASO that binds to the splice acceptor site is called ASO-A, to the donor site, ASO-D and to the site with a variation ASO-M. The ASOs that binds to both the variation sites and either the acceptor or the donor sites are named ASO-MA or ASO-MD respectively. When two ASOs were used in combination the (+) was added between ASO ids. To treat organoids, phosphorodiamidate morpholinos (PMOs) and Endo-Porter peptides with polyethylene glycol conjugates were ordered from Gene Tools, maintaining the same sequence as the ASO tested in patient-derived primary cells. Scramble PMO with 3’-carboxyfluorescein was used to monitor the uptake by various cell types in the organoid.

### Transfection of patient-derived primary cells with ASO

Patient-derived fibroblasts were cultivated and seeded in the 6-well plates for transfection. The RPMI 1640 medium was removed from the plate and washed once with PBS 1X, and Opti-MEM. The transfection solution (Opti-MEM 500µl, Lipofectamine 7μl and 1.5μl of ASO (100μM stock concentration)), consisting of effective ASOs or scramble ASOs, was added to the medium dropwise and incubated for 6h before changing the medium back to high serum regular medium.

### Morpholino treatment in kidney organoids

PMO transfection solution was prepared by mixing 5μM of each PMO with 1ml of BDM and 6μl of Endo-Porter peptide to facilitate efficient cellular uptake. Transfection mix was added to each organoid in a 96-ultra low attachment plate (100μl per well). The treatment protocol lasted for three weeks, with medium changes and transfections every three days continuing until day 38. The study included control groups, such as untreated organoids and organoids treated with labeled-PMO, to evaluate transfection efficiency.

### Transmission electron microscopy

Organoids were fixed in 2.5% glutaraldehyde solution for 2h, washed with 1x PBS three times and stored in 4% PFA in PBS before processing in the Electron Microscopy Platform at Manchester University. Organoids were washed with Hepes buffer (0.1M pH 7.2) several times, then stained with reduced osmium (1.5% potassium ferrocyanide and 1% osmium tetroxide in 0.1M Cacodylate buffer pH 7.2) for 1h and with 1% uranyl acetate in water overnight. They were dehydrated in ethanol or acetone series, embedded into TAAB Low Viscosity resin and polymerized at 6°C for 24h. The 70 nm sections were cut with a Leica UC7 ultramicrotome and placed on a 1x2 mm copper slot grid coated with formvar/carbon film. Images were taken with ThermoFisher Talos L120C electron microscope at 120kV acceleration voltage using Ceta camera.

### Statistics and reproducibility

Statistical analysis for transcript quantification using fragment analysis, RT-qPCR, and macro-enabled protein quantity assessment was conducted using GraphPad Prism (v10.3.0). For two-group comparisons, t-tests were applied to parametric data, whereas multi-group comparisons were analyzed using Two-way ANOVA with Tukey’s post-hoc correction for multiple comparisons. Differentially expressed genes (DEGs) were identified based on an adjusted p-value threshold using the Benjamini-Hochberg method for false discovery rate (FDR) correction. Principal component analysis (PCA), multidimensional scaling (MDS), and unsupervised hierarchical clustering were performed using Rstudio, and plots were generated using the ggplot2 package. For all statistical analyses, significance was defined at an adjusted p-value threshold of <0.05, and details of specific tests (e.g., fold-change cutoffs, threshold settings) are reported in the relevant results sections.

## Supporting information

Supplementary figures and small tables

Supplementary large tables

## List of Supplementary Materials

**Table S1.** Bulk RNA-seq result comparing kidney organoids at day 32 vs day 22 of culture (Excel file)

**Table S2.** Bulk RNA-seq result comparing kidney organoids at day 42 with day 22 of culture (Excel file)

**Table S3.** Protein expression profile in kidney organoids comparing day 34 and day 22 of culture (Excel file)

**Table S4.** Protein expression profile in kidney organoids comparing day 38 and day 22 of culture (Excel file)

**Table S5.** Marker genes obtained from single-cell RNA-seq for annotating different cell-types in kidney organoids (Excel file)

**Table S6.** Differential expressed genes identified in single-cell RNA-seq experiment comparing mutants with isogenic controls (Excel file)

**Table S7.** Pathways and gene ontologies identified from comparing mutants with isogenic controls (Excel file)

## Acknowledgements

We extend our gratitude to addgene for providing the full-length *COL4A5* pNLF1-C_hCOL4A5-Nluc plasmid (#198113), which we used to generate *COL4A5*-specific probes for capturing *COL4A5* cDNA in targeted RNA sequencing. We are also grateful to the Furthmayr lab and the Developmental Studies Hybridoma Bank (DSHB) for supplying the COL4A1/COL4A2 antibody (clone M3F7). We thank Alexander Mironov, the head of the electron microscopy facility at Manchester University for processing and analyzing kidney organoids. We are highly grateful to the Genomics, the Bioinformatics, the Histology and the Cell Imaging platforms of the Structure Fédérative de Recherche (SFR) Necker. We acknowledge the use of the bioresources of the Necker Imagine DNA biobank (BB-033-00065) (patient fibroblasts). This work was supported by funds from the Orphan Kidney Diseases (ORKiD) healthcare network (2021 and 2023, to G.D.), the Agence Nationale de la Recherche under “Investissements d’avenir” program, (ANR-10-IAHU-01) (Institut Imagine) and the “France 2030 program” (DOS0212694) (to C.A. and G.D.). H.S. is a fellow in the Pasteur-Paris University (PPU)-Imagine International Doctoral Program supported by the Institut Imagine.

## Author contributions

H.S, Conceptualization, Data curation, methodology, formal data analysis, Bioinformatic analysis including, single cell data analysis, bulk RNA-seq data analysis, and proteomics data analysis, Imaging and data analysis, primary and stem cell culture and their maintenance and differentiation, resources, visualization, writing the original draft, review and editing; B.S, genome editing in iPSCs, stem cell maintenance and culture, organoid differentiation; N.G, developing the Fiji macro, testing; M.E, stem cell maintenance and culture, organoid differentiation, harvesting and molecular analysis; J.H, stem cell maintenance and culture; O.G, Writing - review and editing; C.A, primary cell maintenance and culture; V.M, targeted RNA sequencing, data curation; P.T, Sample enrichment for proteomics, review and editing; R.L, Funding the proteomics experiments, Writing - review and editing; C.A, Conceptualization, Funding acquisition, Methodology, Writing - review and editing; G.M, Conceptualization, Funding acquisition, Methodology, Writing - review and editing; G.D, Conceptualization, Funding acquisition, Methodology, Writing - original draft, Writing - review and editing

## Data availability

The bulk and single cell RNA-seq data have been deposited to the NCBI GEO with the dataset identifier <u>GSE281080</u> for bulk and <u>GSE281081</u> for the single cell RNA-seq dataset respectively (reviewer token is generated for GSE281080: srgjgggavvqrbgp; and for GSE281081: sbqfkaamjfarfgv). The mass spectrometry proteomics data have been deposited to the ProteomeXchange Consortium via the PRIDE partner repository(70) with the dataset identifier: PXD057360. The Fiji macro for basement membrane protein quantification is provided in the GitHub page (https://github.com/hassansaei/BMQuant).

---

**Fig. S1:**
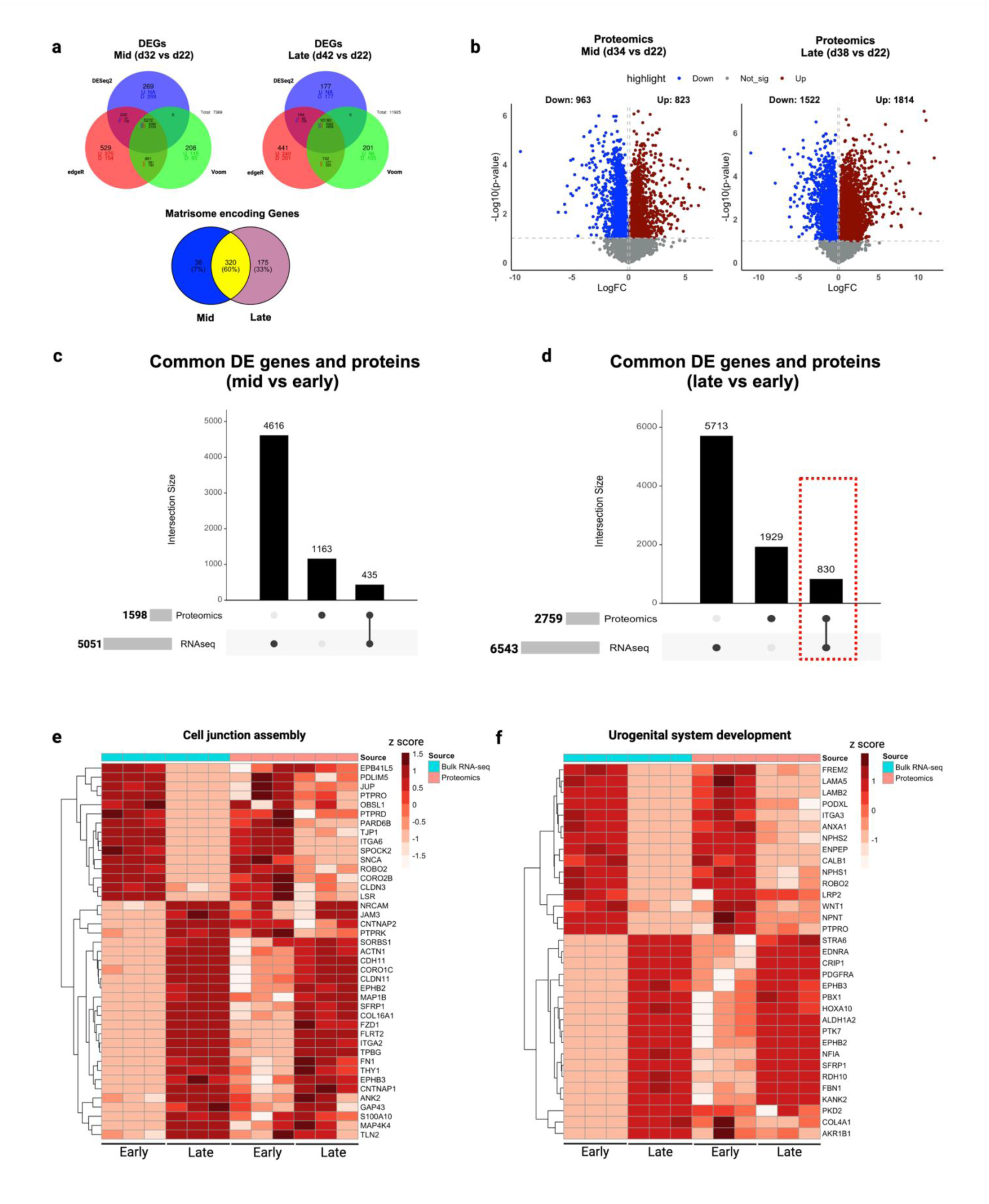
Cross-matching gene and protein expression data between organoids obtained from early and late culture. **(a)** Venn diagrams of DEGs obtained from three different statistical methods (DESeq2, edgeR, and limma-voom, fold change:1.2) at day 32 vs day 22 (mid) and 42 vs day 22 (late). In the mid group, 5,272 DEGs were identified, with 3,089 upregulated and 2,183 downregulated genes. In the late group, 10,180 DEGs were found, with 5,322 upregulated and 4,858 downregulated genes. We used the gene list from the NABA matrisome library (1,025 matrix protein-encoding genes) to identify differentially regulated ECM-encoding genes in each group, with an overlap in matrisome-encoding genes. A total of 320 shared ECM-encoding genes were identified between mid and late condition. Additionally, 175 unique genes were enriched at day 42, indicating the evolution of gene expression beyond day 32. **(b)** Protein groups identified from independent soluble fraction and ECM fraction proteomics analyses were merged and 1786 differentially regulated proteins (d34 vs d22), and 3336 differentially regulated proteins comparing d38 vs d22 (late) were identified. (**c**) and (**d**) Cross matching of RNA and protein data identified 830 shared differentially regulated biomolecules (genes and proteins) comparing late groups. With performing gene set enrichment analysis of these biomolecules, we discovered different pathways and biological processes (refer to Fig. 1e for top 10 in each). **(e)** and **(f)** Heatmaps of genes and proteins enriched in GO terms related to urogenital system development and cell junction assembly, respectively. These terms were enriched following focused analysis of shared differentially expressed genes and proteins from late group comparison.

**Fig. S2:**
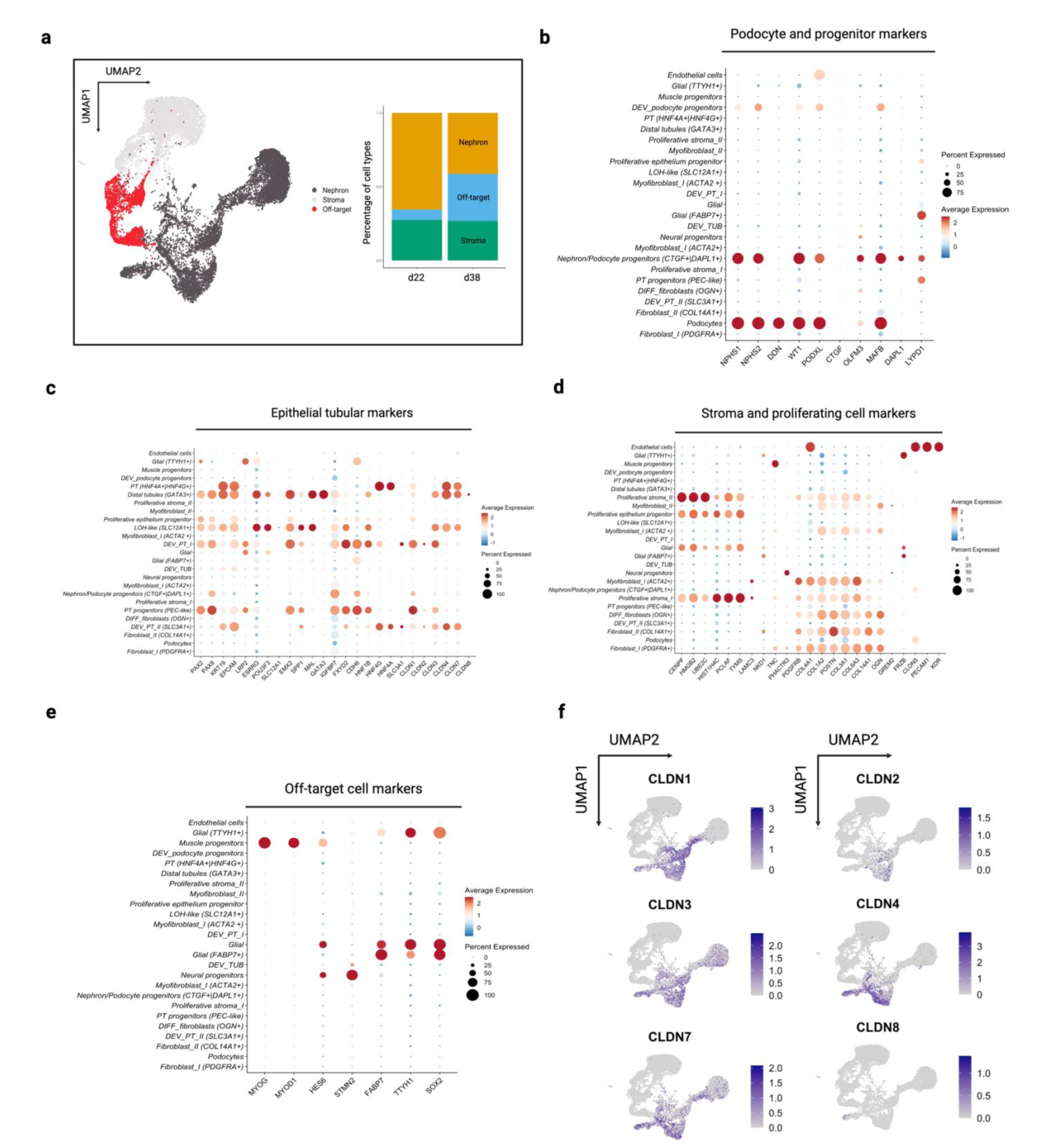
Single-cell RNA profiling of kidney organoids identifies nephron, stromal, and off-target cell populations. **(a)** UMAP plot highlighting nephron, stromal, and off-target cell populations with a corresponding bar plot showing the proportional distribution of each population within the organoid. **(b)** Dot plot showing the expression of late podocyte and podocyte progenitor markers across different cell types, highlighting the specificity of markers like *NPHS1, NPHS2, PODXL, DDN*, and *WT1* in podocytes. **(c)** Dot plot of tubular epithelial markers, demonstrating distinct expression patterns across various nephron segments, including proximal *PAX2, PAX8, EPCAM*, proximal tubule markers (*CDH6, SLC3A1, HNF4G* and *HNF4A*) and distal tubule (*GATA3*, and *MAL*), LoH-like cells expressing *POU3F3*, and *ESRRG*. **(d)** Dot plot illustrating stromal and proliferating cell markers, with prominent expression of genes like *COL1A1* and *ACTA2* in stromal clusters (myofibroblasts), and open chromatin markers (*PCLAF*, and *TYMS*) in proliferating cells. **(e)** Dot plot of off-target cell markers, showing expression of neural progenitor (*STMN2*, and *HES6*) and muscle progenitor markers (*MYOG*, and *MYOD1*). **(f)** Feature plots showing expression of claudin family genes (*CLDN1* to *CLDN8*) across the organoid, indicating their segment-specific enrichment in nephron and tubular epithelial clusters (*CLDN1* in PEC-like cells and *CLDN4* more in distal part of the nephron).

**Fig. S3:**
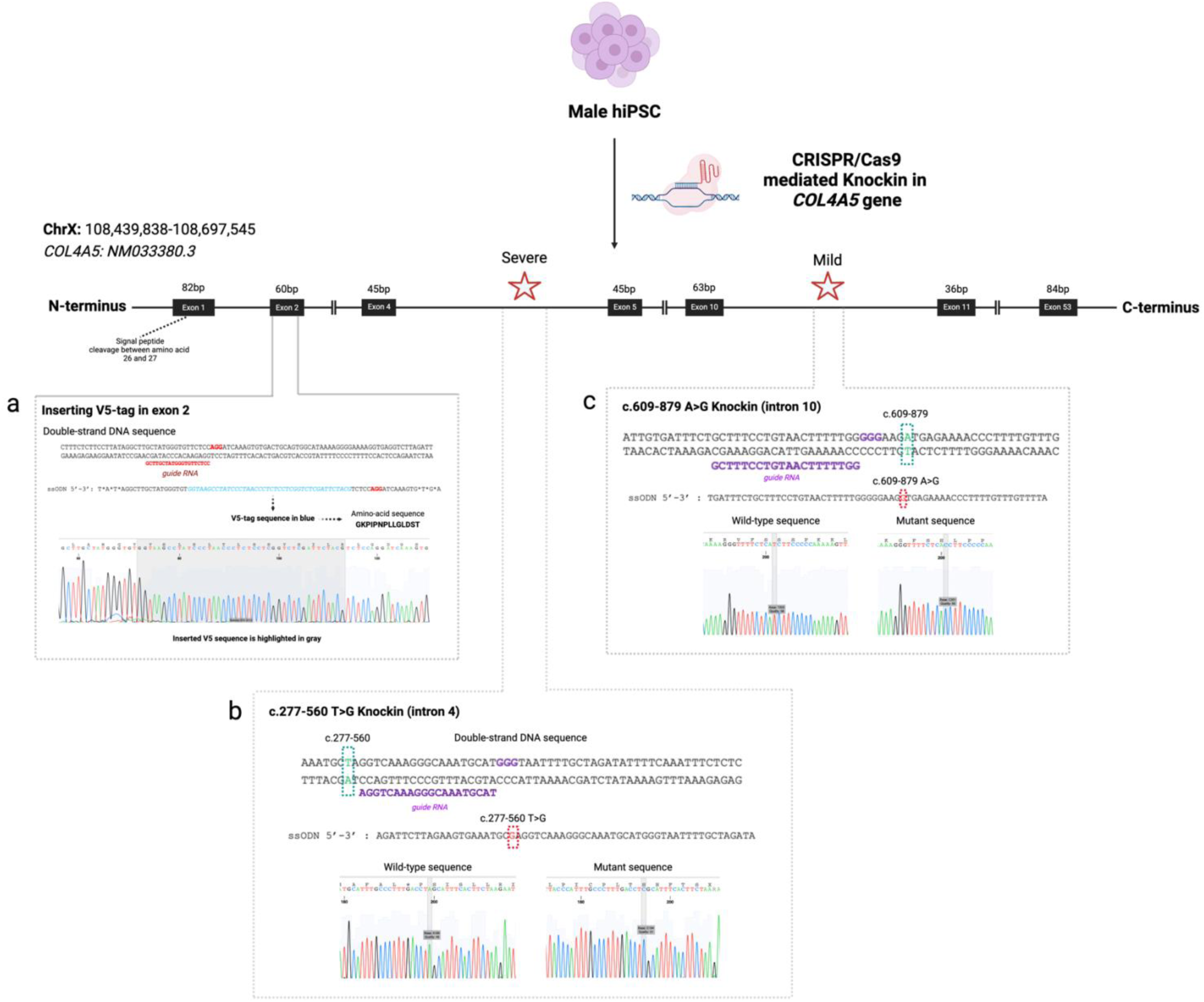
Generation of isogenic hiPSC lines with CRISPR/Cas9-mediated knock-in of *COL4A5* variants. The figure illustrates the CRISPR/Cas9 strategy used to introduce V5-tag **(a)**, severe **(b)** and mild **(c)** *COL4A5* variants in male hiPSCs for modeling X-linked Alport Syndrome. The top panel provides an overview of the *COL4A5* gene structure, highlighting the locations of the introduced variants. The left inset details the V5-tag knock-in strategy in exon 2, showing the insertion site (gray highlighted sequence) with Sanger sequencing confirming successful integration. The bottom-left inset depicts the severe c.277-560T>G variant knock-in in intron 4, showing the guide RNA (purple) and donor ssODN used for editing, along with Sanger sequencing verifying the mutation. The right inset outlines the mild c.609+879A>G variant knock-in in intron 10, presenting the guide RNA (purple), donor ssODN, and corresponding Sanger sequencing chromatograms confirming the mutant sequence. The generated clones were checked for karyotype and genome instability using SNP array.

**Fig. S4:**
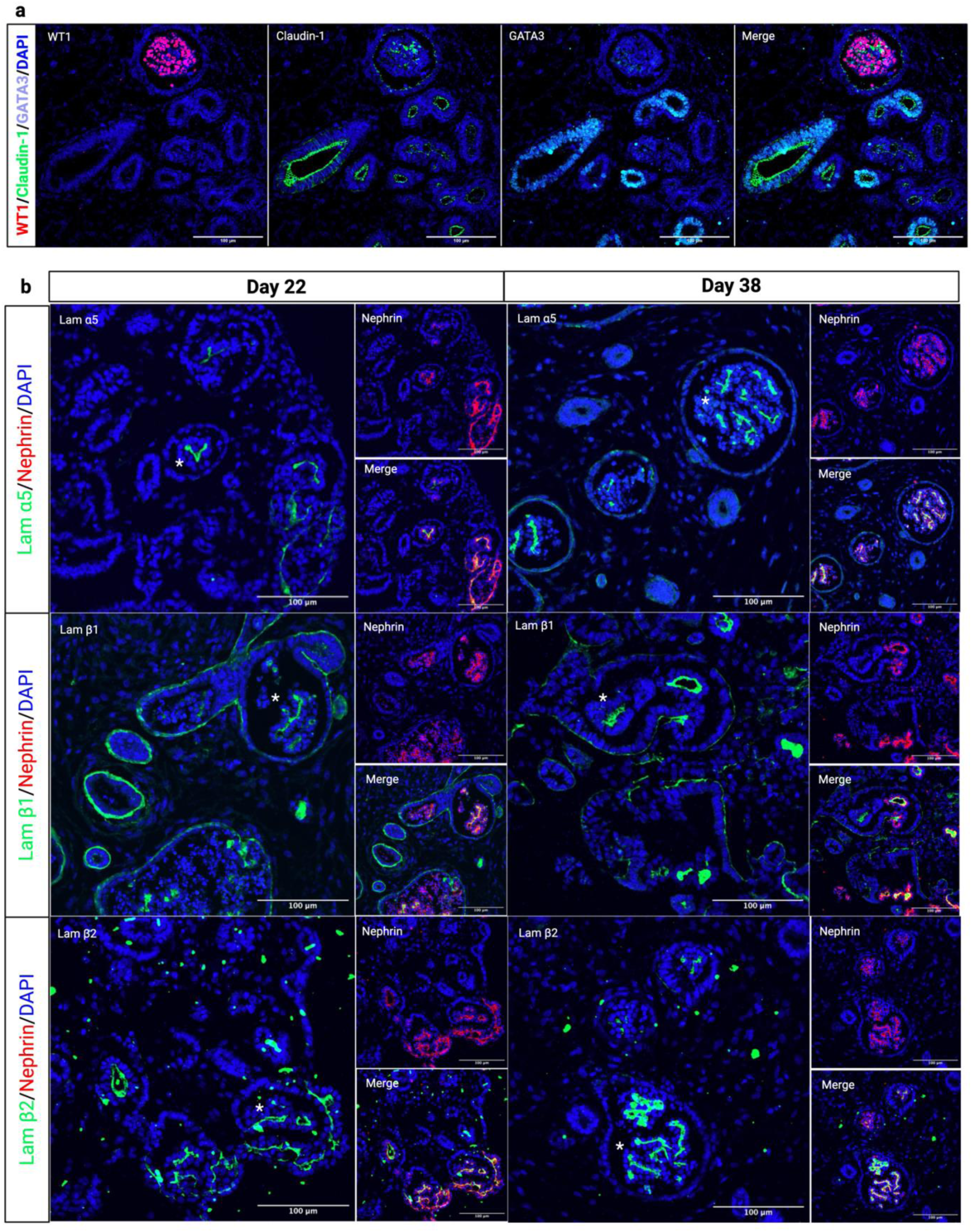
Immunofluorescence staining of kidney organoids highlighting the parietal epithelial cells and the localization of different laminins in the BMs. **(a)** Spatial distribution of WT1 (in red), claudin-1 (in green), and GATA Binding Protein 3 (in cyan) in organoid (day 38). As expected, WT1 is mainly localized to the nucleus of the podocytes, while claudin-1 is present in the parietal epithelial-like cells (PEC-like) and distal tubules (GATA3+). These cells expressing collagen type IV network and laminin proteins. Panels **(b)** compared laminin α5, β1, and β2 localization and abundance comparing days 22 and 38 of organoid culture (see * for the glomeruli). This panel confirming the presence of laminin mature network in organoids early in the organoid development.

**Fig. S5:**
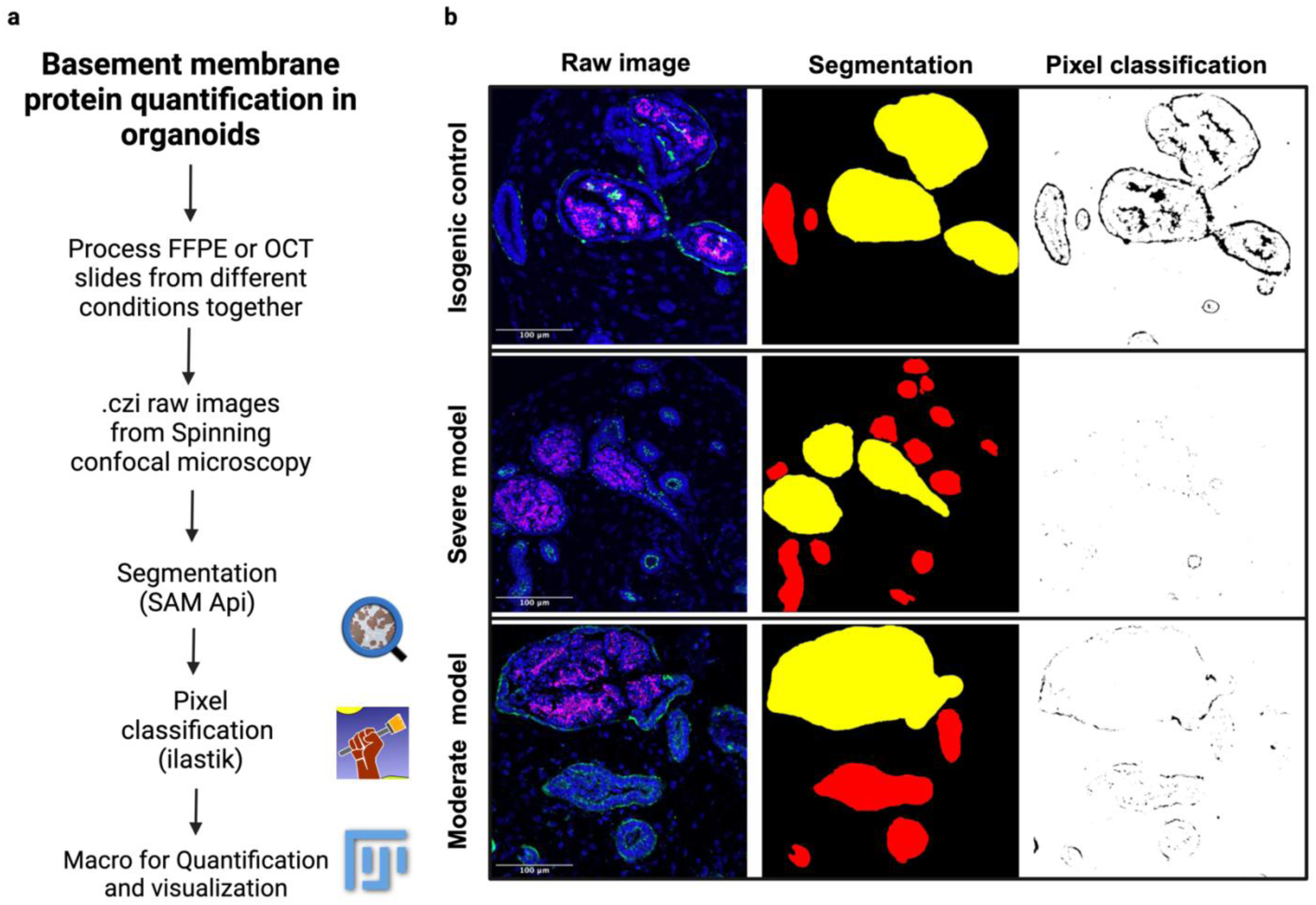
Development of a Fiji macro for quantification of basement membrane protein expression in kidney organoids. **(a)** The workflow begins with the segmentation of glomeruli and tubules using QuPath, followed by pixel classification with ilastik to detect true signals. SAM API was utilized for accurate identification of object borders, and the ROIs were saved for use in the macro. Each fluorescent channel corresponding to a specific protein was then seperated. The channel of interest was imported into ilastik for random forest model-based pixel classification. Using the QuPath and ilastik project files, the macro scans the raw images and performs quantification, generating CSV files that contain measurements such as the full border area, mean channel intensity in the border and inside the object, and channel area. The quantity of protein in the BM of tubules (TBM) and glomeruli (PEC-BM) was calculated by multiplying the mean channel intensity in the border by the border area or the mean channel intensity inside the glomeruli multiply by the area inside the glomeruli (GBM). This method allows us to measure the quantity of collagen α5(IV) protein in the BMs of the isogenic control, severe, and moderate models before and after ASO treatment.

**Fig. S6:**
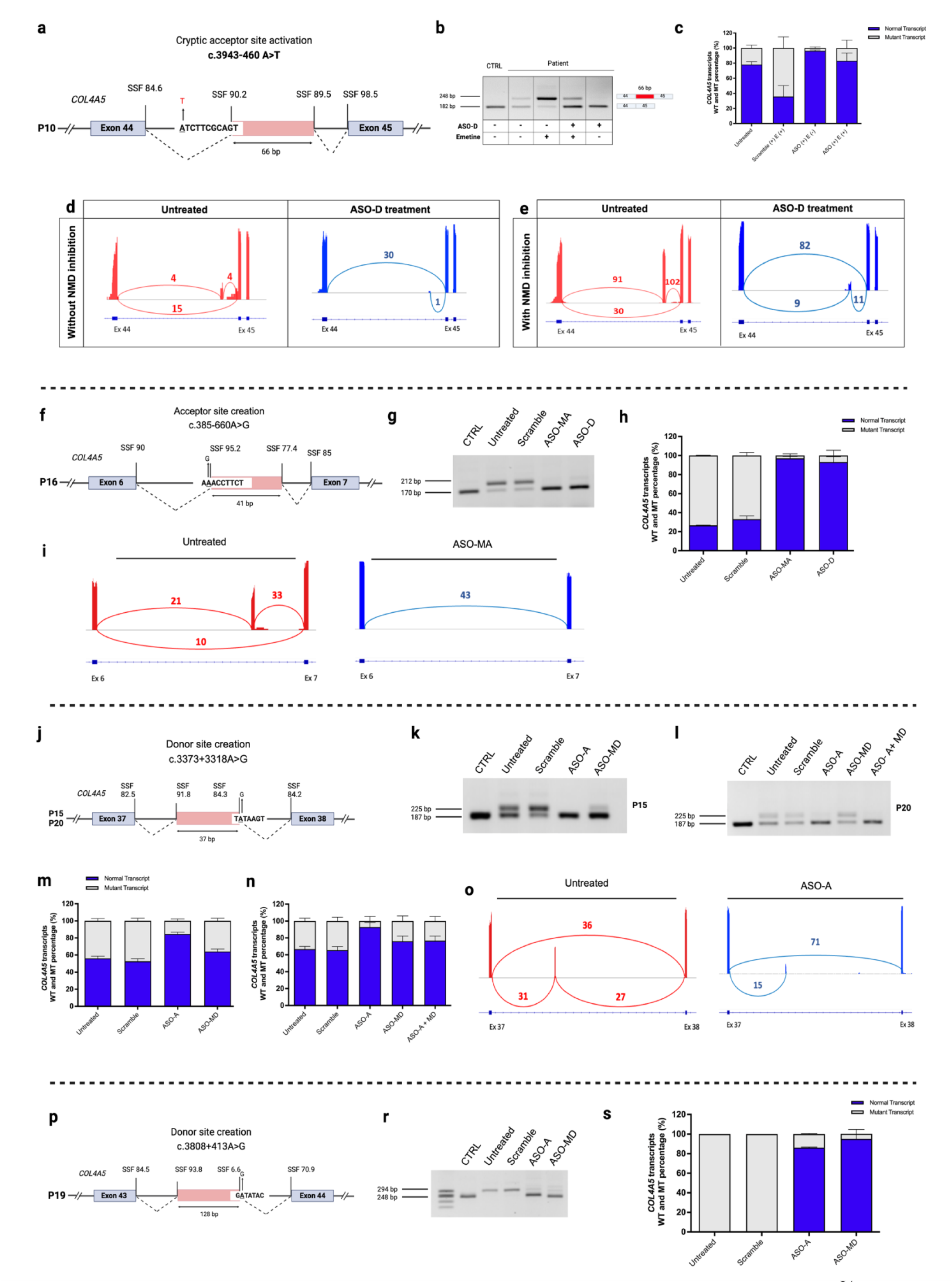
Antisense oligonucleotides targeting cryptic splice sites in multiple *COL4A5* deep-intronic variants showed efficient splicing correction. **(a)** P10 variant activated cryptic acceptor site at intron 44. **(b)** RT-PCR results confirmed aberrant splicing in the patient cells with NMD-mediated mRNA degradation, with ASO-D treatment restoring normal splicing. **(c)** Quantification of wild-type and mutant transcripts confirmed splice correction after ASO treatment. Panels **(d)** and **(e)** showing splice junction usage with and without NMD inhibition, and successful pseudo-exon skipping after ASO treatment. **(f)** P16 variant created novel acceptor site resulting in 41bp retention of intron 6 sequence in the *COL4A5* mRNA **(g)**. This aberrant splicing event was successfully corrected following ASO-MA and ASO-D treatment. Panels **(h)** and **(i)** showed the quantification of *COL4A5* transcripts and splice junction usage before and after ASO treatment. **(j)** In P15 and P20 the variant created a novel donor site, and RT-PCR analysis **(k)** showed the retention of 37bp from intron 37 which was effectively restored by ASO-A treatment. This was further validated by fragment analysis **(m, and n)** and targeted RNA-seq **(o). (p)** Details of a donor site creation in P19. RT-PCR analysis **(r)** demonstrated the restoration of normal transcript levels following ASO-A and ASO-MD treatment. **(s)** The effects of ASO treatments on transcript levels, were further quantified by fragment analysis.

**Table S8.**
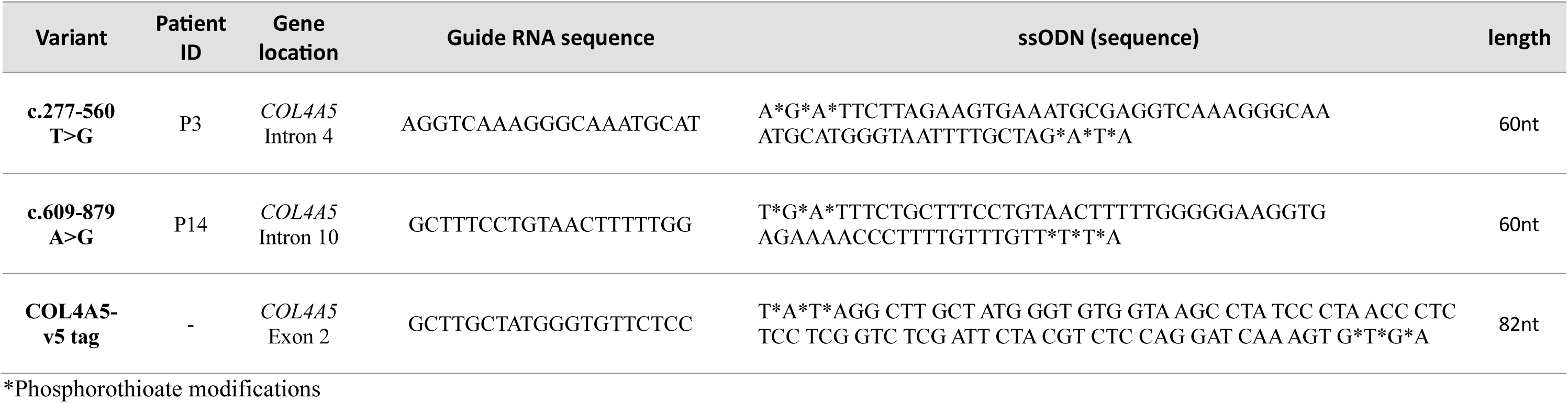
Guide RNAs and ssODN used for CRISPR/Cas9 mediated knock-ins.

**Table S9.**
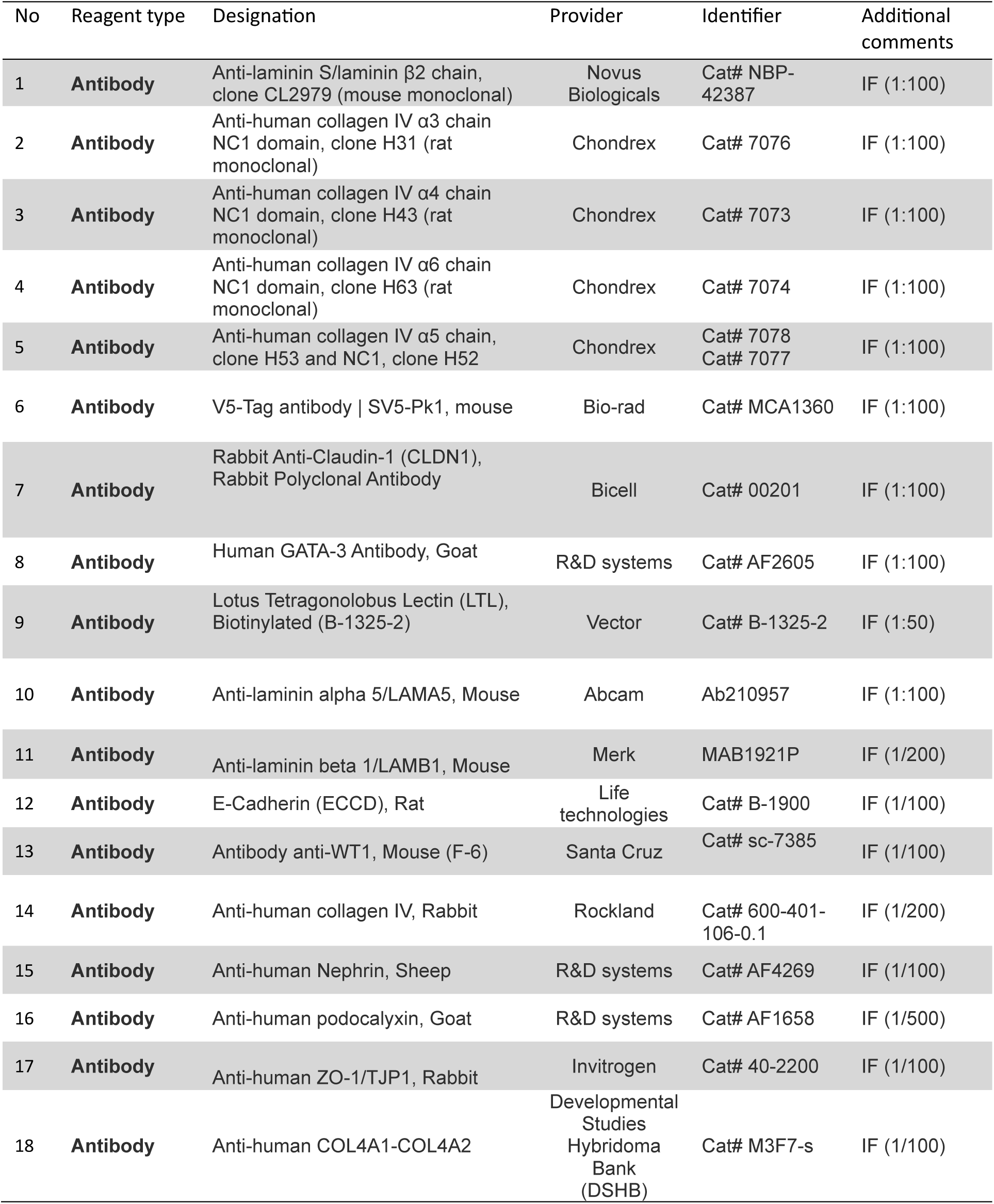
List of the antibodies used in this study.

**Table S10.**
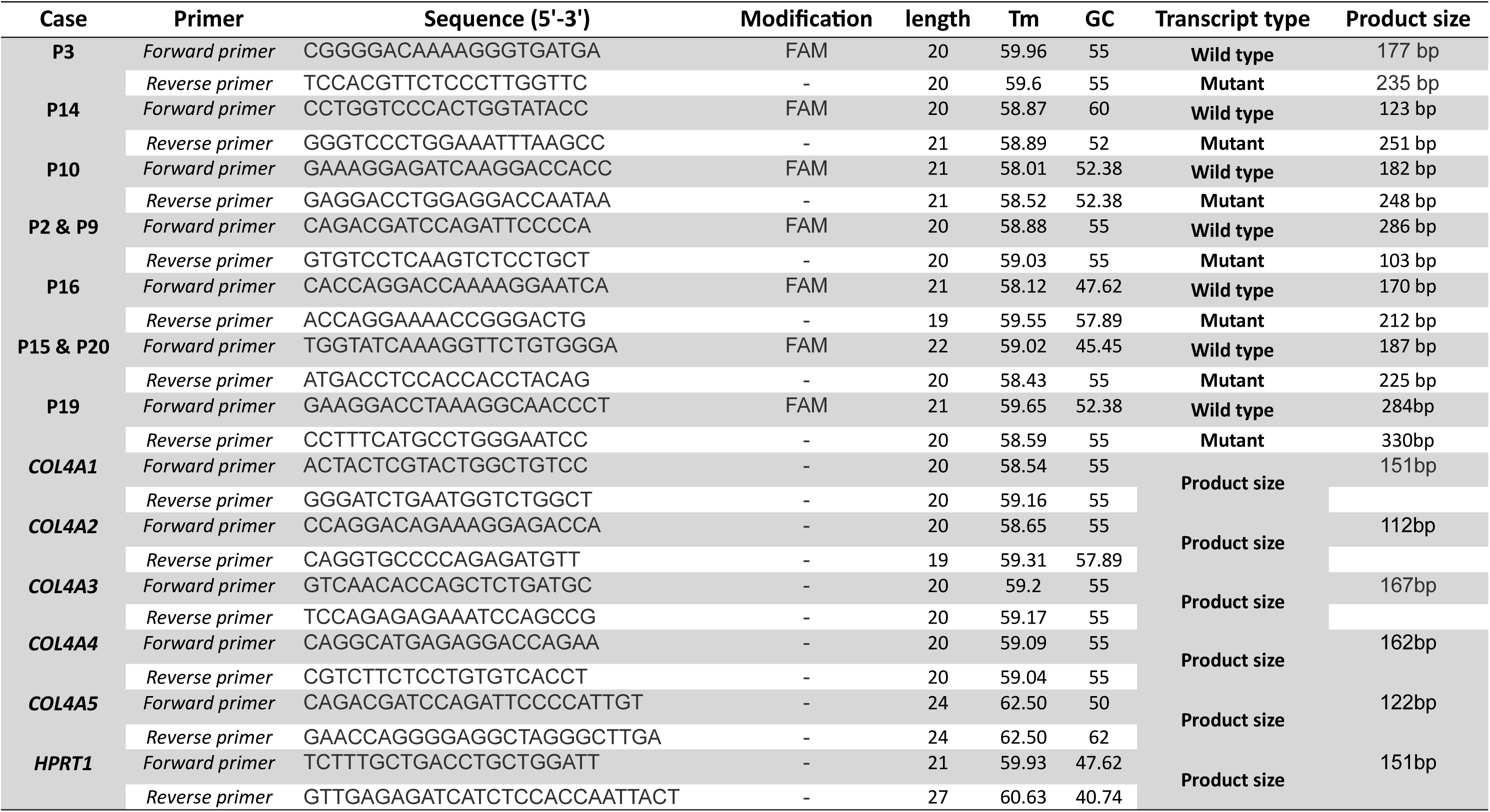
Primers used for RT-PCR and RT-qPCR.

