## Supplementary figures and small tables for "Therapeutic splice modulation of COL4A5 reinstates collagen IV assembly in an organoid model of X-linked Alport syndrome"

**Supplementary information**

**Supplementary Figures:**

**Fig. S1:** Cross-matching gene and protein expression data between organoids obtained from early and late culture.

**Fig. S2:** Single-cell RNA profiling of kidney organoids identifies nephron, stromal, and off-target cell populations.

**Fig. S3:** Generation of isogenic hiPSC lines with CRISPR/Cas9-mediated knock-in of *COL4A5* variants

**Fig. S4:** Immunofluorescence staining of kidney organoids highlighting the parietal epithelial cells and the localization of different laminins in the BMs

**Fig. S5:** Development of a Fiji macro for quantification of basement membrane protein expression in kidney organoids

**Fig. S6:** Antisense oligonucleotides targeting cryptic splice sites in multiple *COL4A5* deep-intronic variants showed efficient splicing correction

**Supplementary Tables:**

**Table S1.** Bulk RNA-seq result comparing kidney organoids at day 32 vs day 22 of culture

**Table S2.** Bulk RNA-seq result comparing kidney organoids at day 42 with day 22 of culture

**Table S3.** Protein expression profile in kidney organoids comparing day 34 and day 22 of culture

**Table S4.** Protein expression profile in kidney organoids comparing day 38 and day 22 of culture

**Table S5.** Marker genes obtained from single-cell RNA-seq for annotating different cell-types in kidney organoids

**Table S6.** Differential expressed genes identified in single-cell RNA-seq experiment comparing mutants with isogenic controls

**Table S7.** Pathways and gene ontologies identified from comparing mutants with isogenic controls

**Table S8.** Guide RNAs and ssODN used for CRISPR/Cas9 mediated knockins

**Table S9.** List of the antibodies used in this study

**Table S10.** Primers used for RT-PCR and RT-qPCR

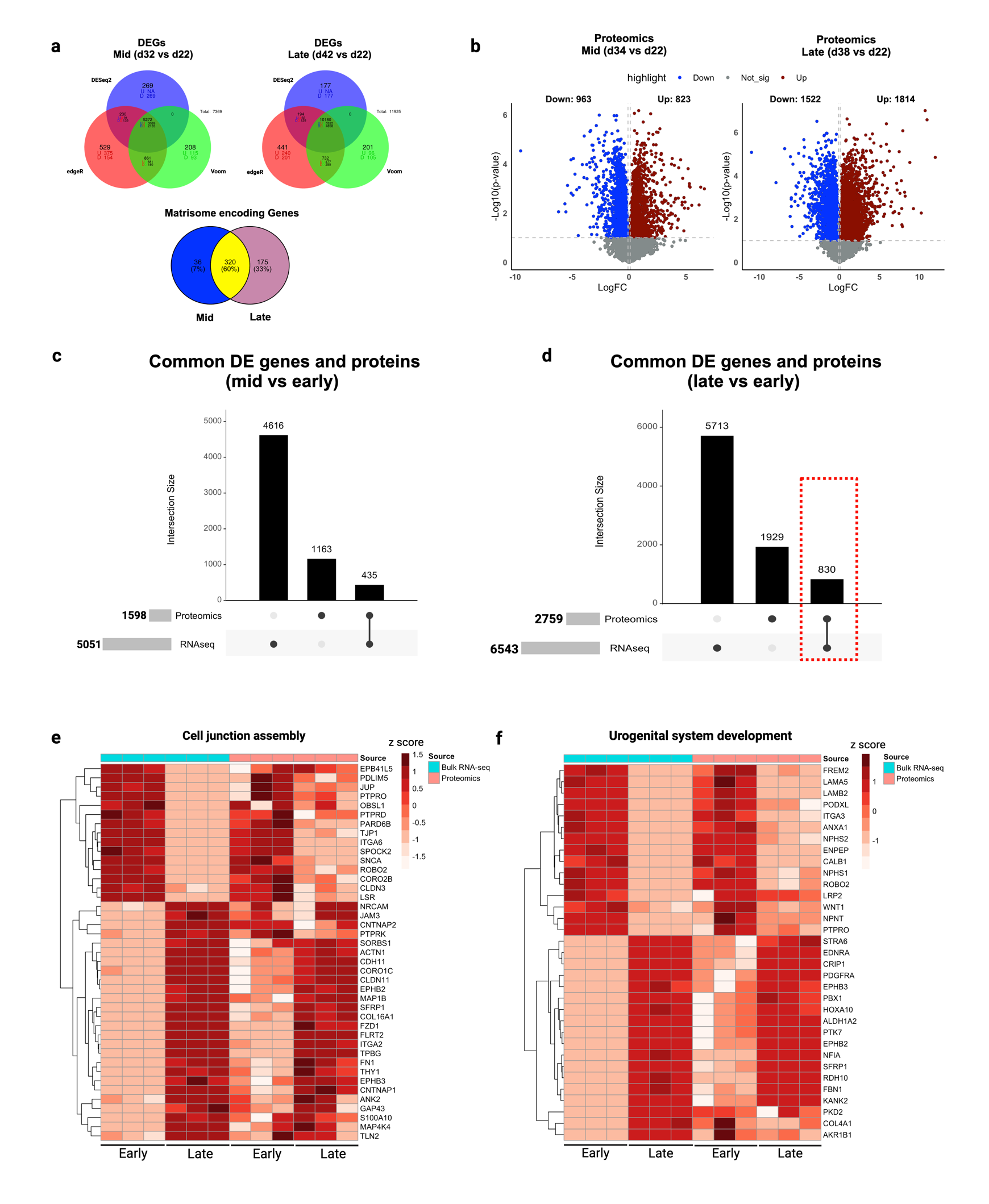

**Fig. S1: Cross-matching gene and protein expression data between organoids obtained from early and late culture**

**(a)** Venn diagrams of DEGs obtained from three different statistical methods (DESeq2, edgeR, and limma-voom, fold change:1.2) at day 32 vs day 22 (mid) and 42 vs day 22 (late). In the mid group, 5,272 DEGs were identified, with 3,089 upregulated and 2,183 downregulated genes. In the late group, 10,180 DEGs were found, with 5,322 upregulated and 4,858 downregulated genes. We used the gene list from the NABA matrisome library (1,025 matrix protein-encoding genes) to identify differentially regulated ECM-encoding genes in each group, with an overlap in matrisome-encoding genes. A total of 320 shared ECM-encoding genes were identified between mid and late condition. Additionally, 175 unique genes were enriched at day 42, indicating the evolution of gene expression beyond day 32. **(b)** Protein groups identified from independent soluble fraction and ECM fraction proteomics analyses were merged and 1786 differentially regulated proteins (d34 vs d22), and 3336 differentially regulated proteins comparing d38 vs d22 (late) were identified. (**c**) and (**d**) Cross matching of RNA and protein data identified 830 shared differentially regulated biomolecules (genes and proteins) comparing late groups. With performing gene set enrichment analysis of these biomolecules, we discovered different pathways and biological processes (refer to Fig. 1e for top 10 in each). **(e)** and **(f)** Heatmaps of genes and proteins enriched in GO terms related to urogenital system development and cell junction assembly, respectively. These terms were enriched following focused analysis of shared differentially expressed genes and proteins from late group comparison.

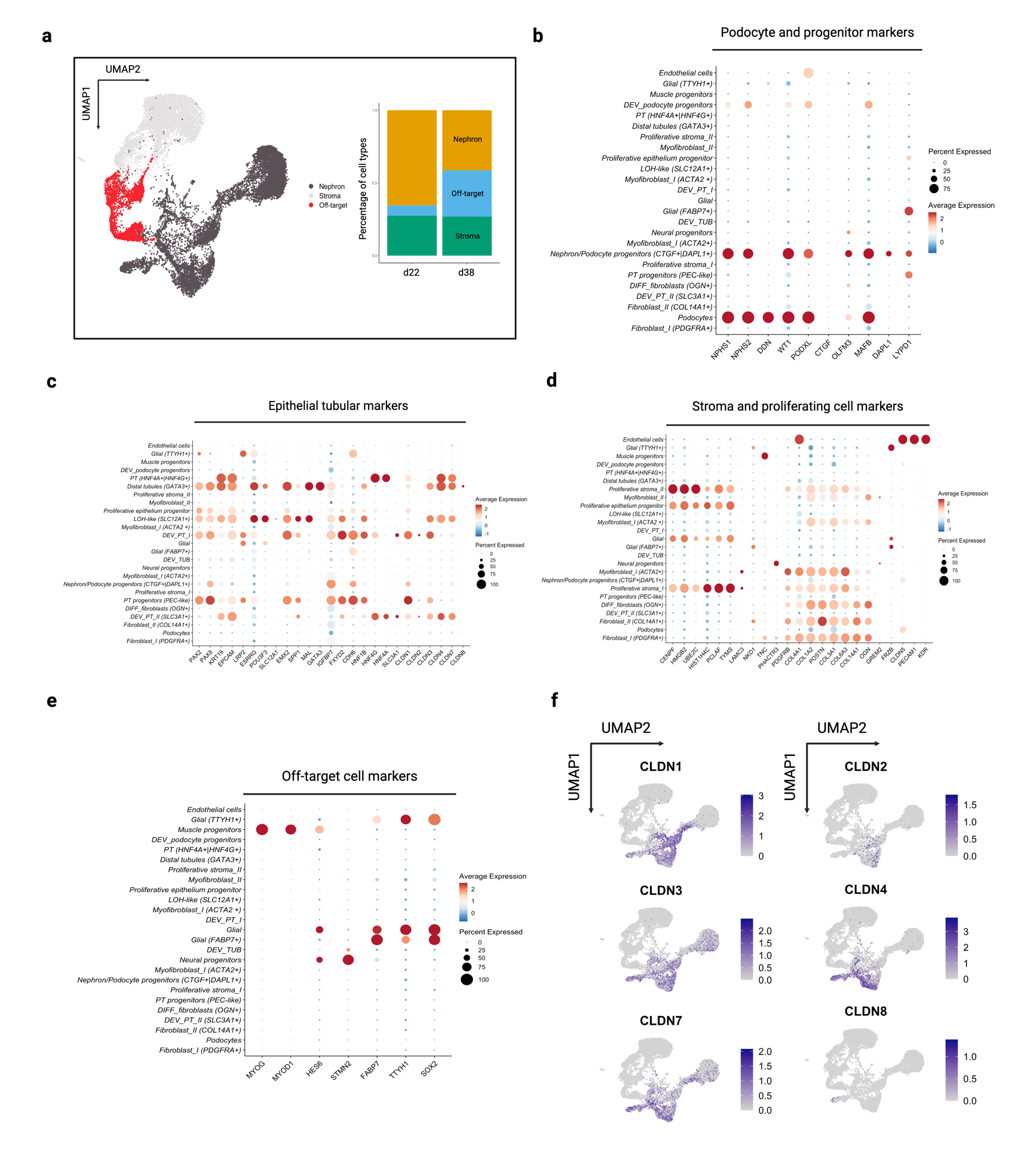

**Fig. S2: Single-cell RNA profiling of kidney organoids identifies nephron, stromal, and off-target cell populations**

**(a)** UMAP plot highlighting nephron, stromal, and off-target cell populations with a corresponding bar plot showing the proportional distribution of each population within the organoid. **(b)** Dot plot showing the expression of late podocyte and podocyte progenitor markers across different cell types, highlighting the specificity of markers like *NPHS1, NPHS2, PODXL, DDN*, and *WT1* in podocytes. **(c)** Dot plot of tubular epithelial markers, demonstrating distinct expression patterns across various nephron segments, including proximal *PAX2, PAX8, EPCAM*, proximal tubule markers (*CDH6, SLC3A1, HNF4G* and *HNF4A*) and distal tubule (*GATA3*, and *MAL*), LoH-like cells expressing *POU3F3*, and *ESRRG*. **(d)** Dot plot illustrating stromal and proliferating cell markers, with prominent expression of genes like *COL1A1* and *ACTA2* in stromal clusters (myofibroblasts), and open chromatin markers (*PCLAF*, and *TYMS*) in proliferating cells. **(e)** Dot plot of off-target cell markers, showing expression of neural progenitor (*STMN2*, and *HES6*) and muscle progenitor markers (*MYOG*, and *MYOD1*). **(f)** Feature plots showing expression of claudin family genes (*CLDN1* to *CLDN8*) across the organoid, indicating their segment-specific enrichment in nephron and tubular epithelial clusters (*CLDN1* in PEC-like cells and *CLDN4* more in distal part of the nephron).

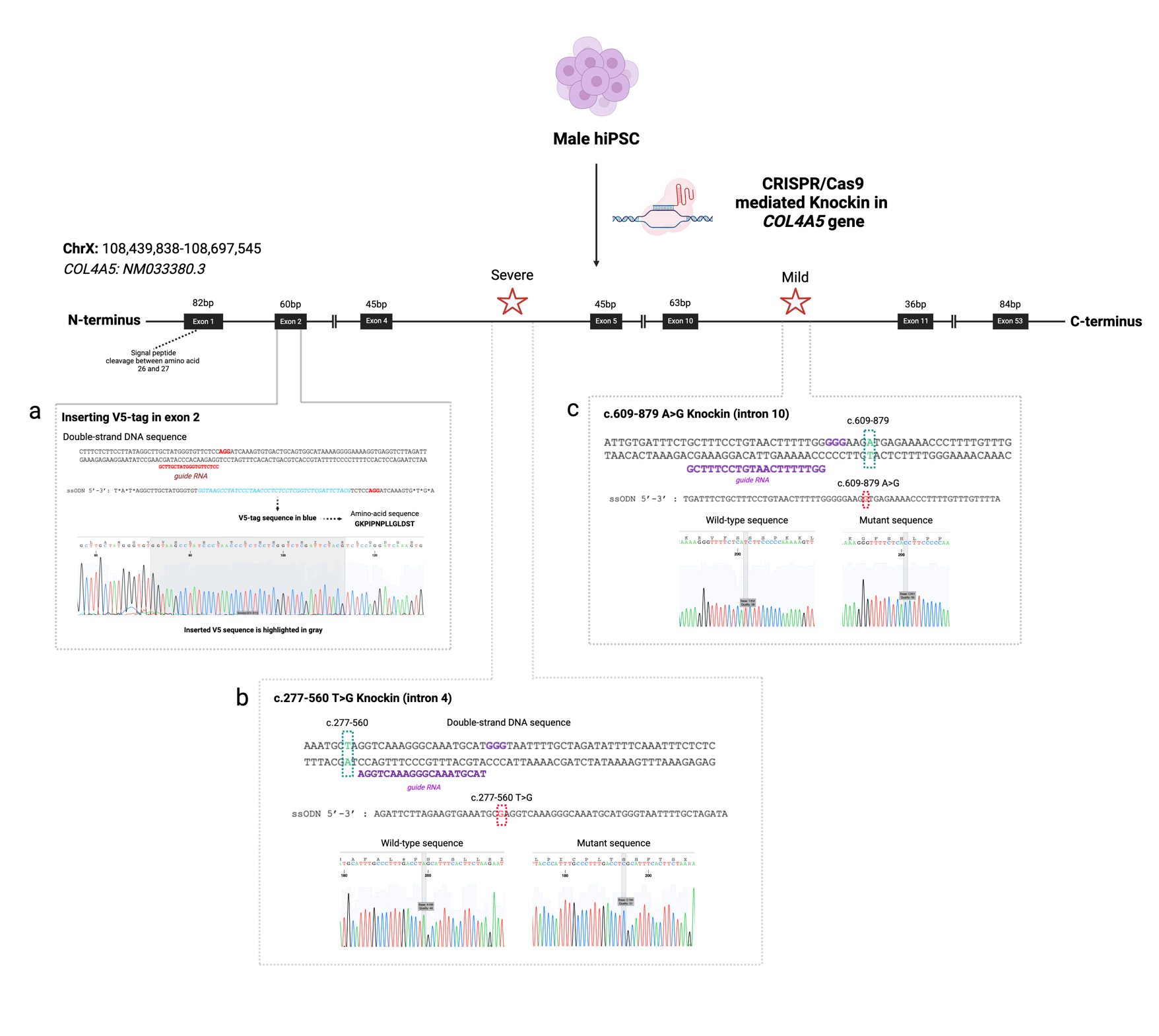

**Fig. S3: Generation of isogenic hiPSC lines with CRISPR/Cas9-mediated knock-in of *COL4A5* variants**

The figure illustrates the CRISPR/Cas9 strategy used to introduce V5-tag **(a)**, severe **(b)** and mild **(c)** *COL4A5* variants in male hiPSCs for modeling X-linked Alport Syndrome. The top panel provides an overview of the *COL4A5* gene structure, highlighting the locations of the introduced variants. The left inset details the V5-tag knock-in strategy in exon 2, showing the insertion site (gray highlighted sequence) with Sanger sequencing confirming successful integration. The bottom-left inset depicts the severe c.277-560T>G variant knock-in in intron 4, showing the guide RNA (purple) and donor ssODN used for editing, along with Sanger sequencing verifying the mutation. The right inset outlines the mild c.609+879A>G variant knock-in in intron 10, presenting the guide RNA (purple), donor ssODN, and corresponding Sanger sequencing chromatograms confirming the mutant sequence. The generated clones were checked for karyotype and genome instability using SNP array.

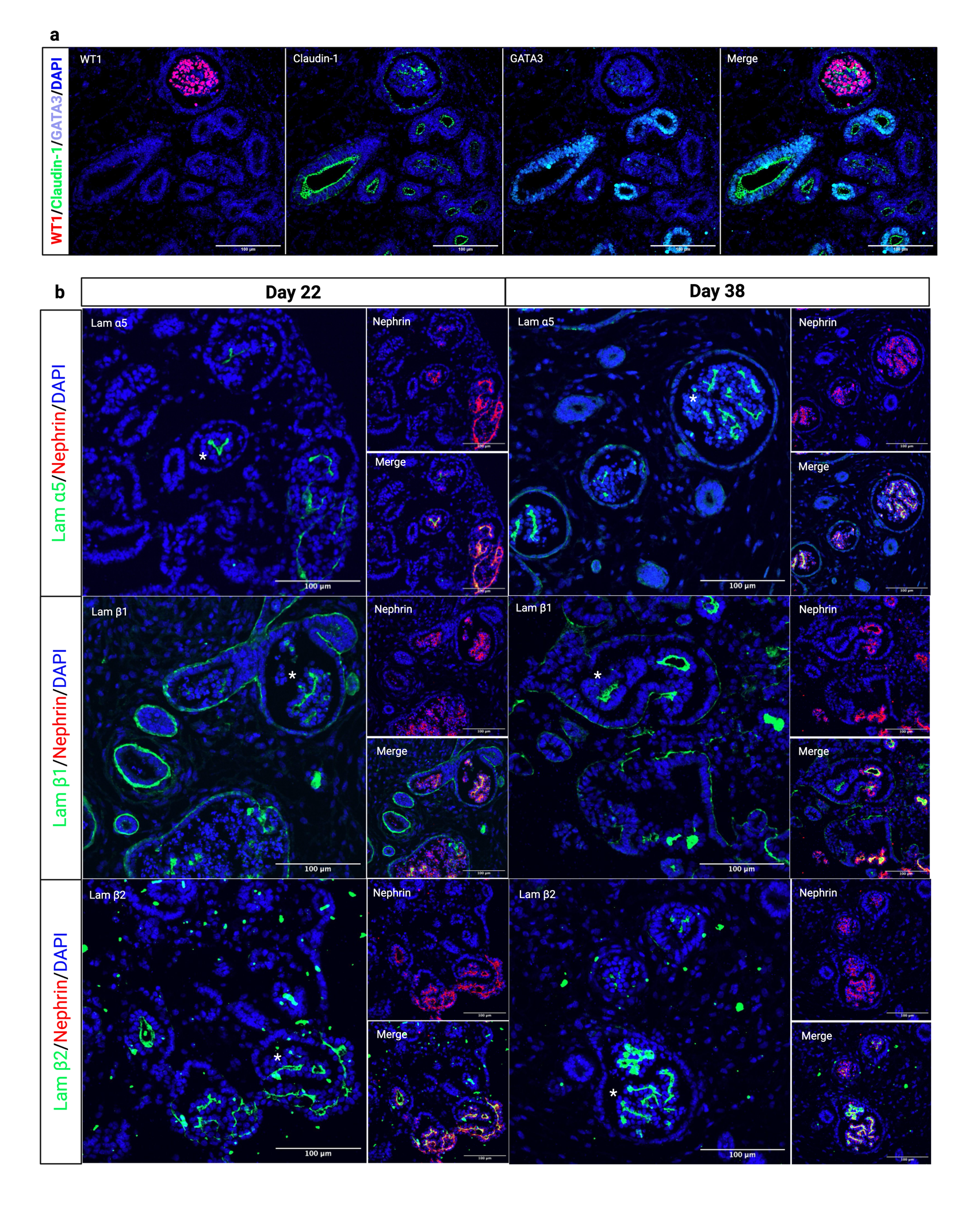

**Fig. S4:** **Immunofluorescence staining of kidney organoids highlighting the parietal epithelial cells and the localization of different laminins in the BMs**

**(a)** Spatial distribution of WT1 (in red), claudin-1 (in green), and GATA Binding Protein 3 (in cyan) in organoid (day 38). As expected, WT1 is mainly localized to the nucleus of the podocytes, while claudin-1 is present in the parietal epithelial-like cells (PEC-like) and distal tubules (GATA3+). These cells expressing collagen type IV network and laminin proteins. Panels **(b)** compared laminin α5, β1, and β2 localization and abundance comparing days 22 and 38 of organoid culture (see * for the glomeruli). This panel confirming the presence of laminin mature network in organoids early in the organoid development.

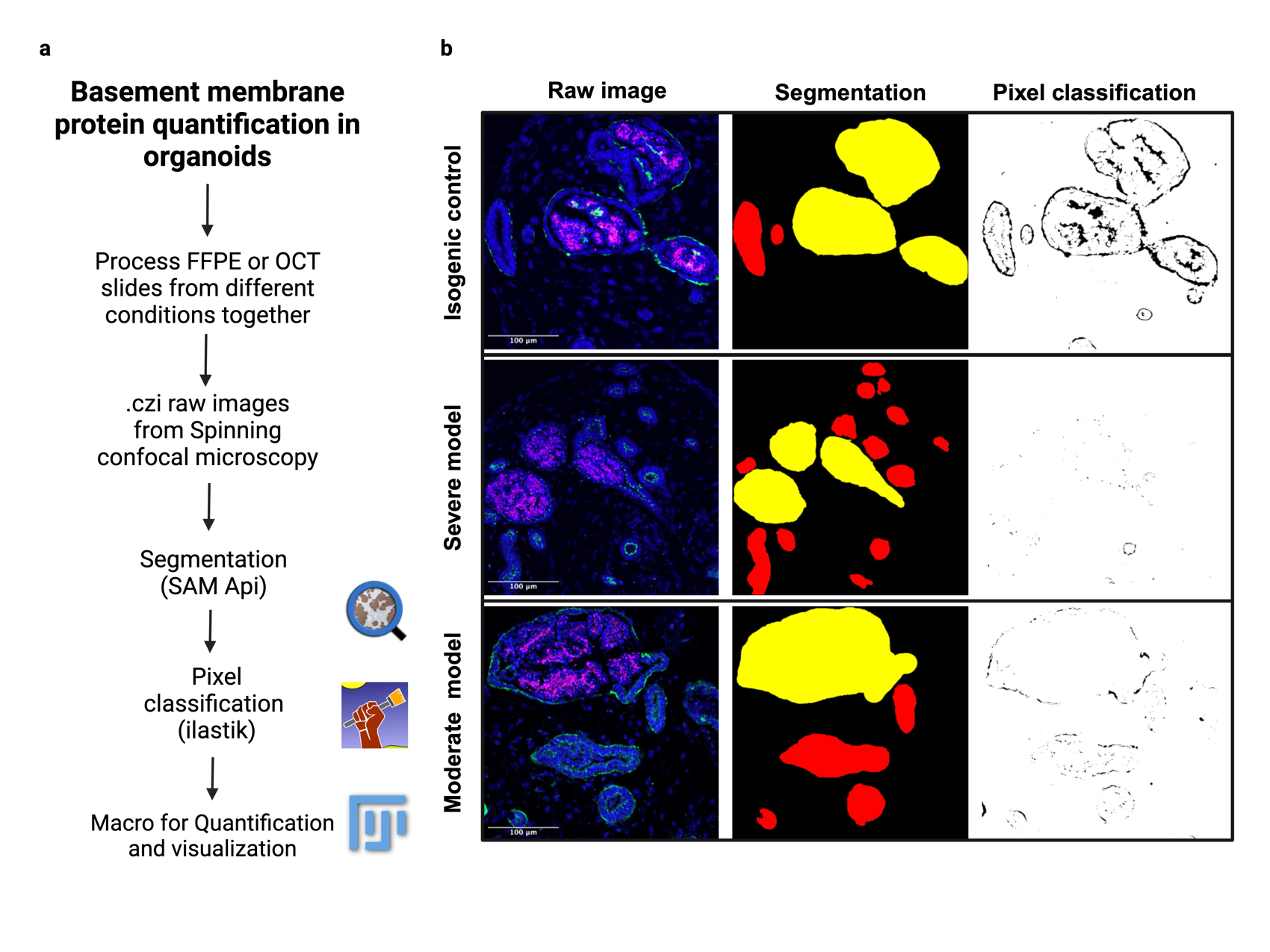

**Fig. S5: Development of a Fiji macro for quantification of basement membrane protein expression in kidney organoids**

**(a)** The workflow begins with the segmentation of glomeruli and tubules using QuPath, followed by pixel classification with ilastik to detect true signals. SAM API was utilized for accurate identification of object borders, and the ROIs were saved for use in the macro. Each fluorescent channel corresponding to a specific protein was then seperated. The channel of interest was imported into ilastik for random forest model-based pixel classification. Using the QuPath and ilastik project files, the macro scans the raw images and performs quantification, generating CSV files that contain measurements such as the full border area, mean channel intensity in the border and inside the object, and channel area. The quantity of protein in the BM of tubules (TBM) and glomeruli (PEC-BM) was calculated by multiplying the mean channel intensity in the border by the border area or the mean channel intensity inside the glomeruli multiply by the area inside the glomeruli (GBM). This method allows us to measure the quantity of collagen α5(IV) protein in the BMs of the isogenic control, severe, and moderate models before and after ASO treatment.

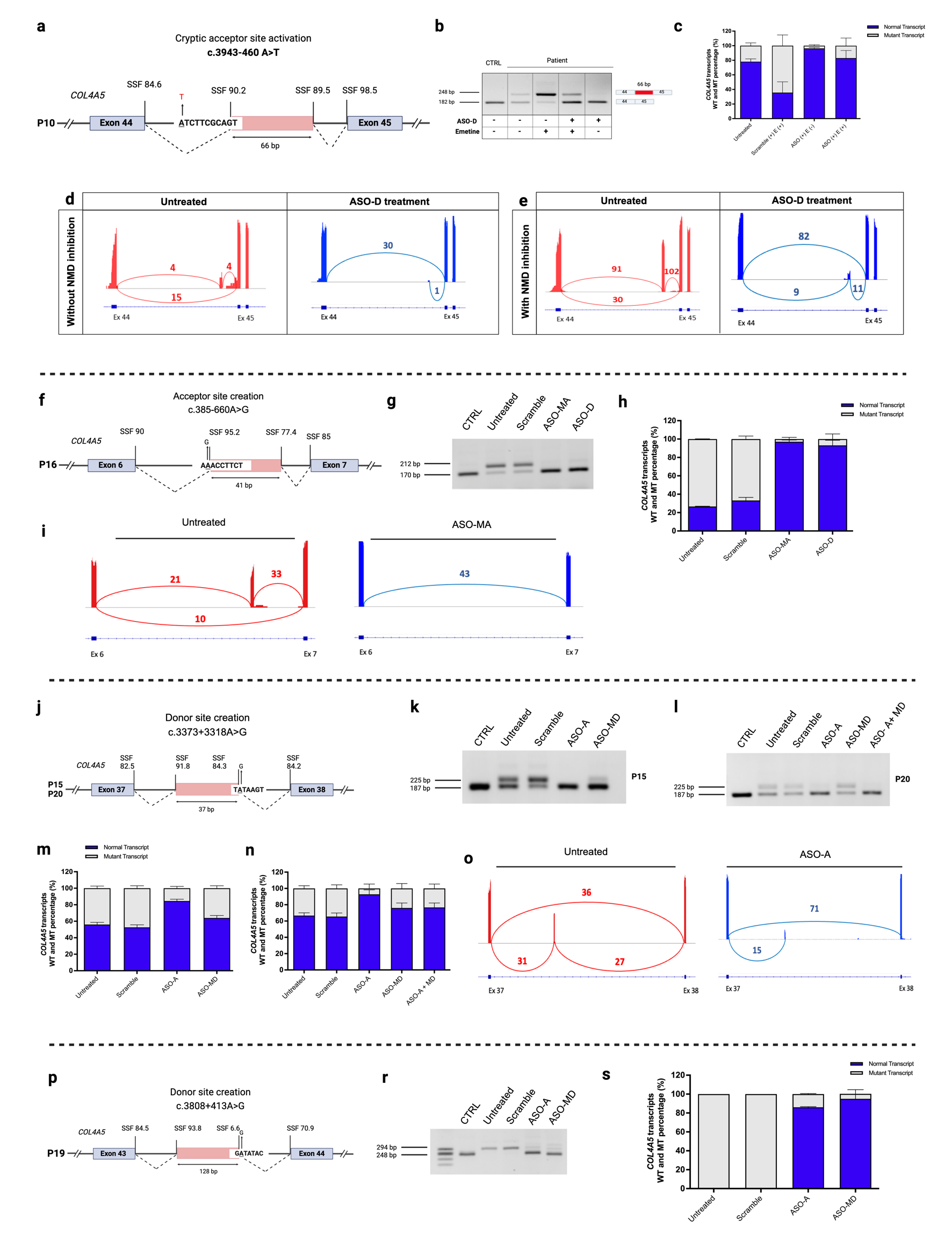

**Fig. S6: Antisense oligonucleotides targeting cryptic splice sites in multiple *COL4A5* deep-intronic variants showed efficient splicing correction**

**(a)** P10 variant activated cryptic acceptor site at intron 44. **(b)** RT-PCR results confirmed aberrant splicing in the patient cells with NMD-mediated mRNA degradation, with ASO-D treatment restoring normal splicing. **(c)** Quantification of wild-type and mutant transcripts confirmed splice correction after ASO treatment. Panels **(d)** and **(e)** showing splice junction usage with and without NMD inhibition, and successful pseudo-exon skipping after ASO treatment. **(f)** P16 variant created novel acceptor site resulting in 41bp retention of intron 6 sequence in the *COL4A5* mRNA **(g)**. This aberrant splicing event was successfully corrected following ASO-MA and ASO-D treatment. Panels **(h)** and **(i)** showed the quantification of *COL4A5* transcripts and splice junction usage before and after ASO treatment. **(j)** In P15 and P20 the variant created a novel donor site, and RT-PCR analysis **(k)** showed the retention of 37bp from intron 37 which was effectively restored by ASO-A treatment. This was further validated by fragment analysis **(m, and n)** and targeted RNA-seq **(o). (p)** Details of a donor site creation in P19. RT-PCR analysis **(r)** demonstrated the restoration of normal transcript levels following ASO-A and ASO-MD treatment. **(s)** The effects of ASO treatments on transcript levels, were further quantified by fragment analysis.

**Table S8. Guide RNAs and ssODN used for CRISPR/Cas9 mediated knock-ins.**

| Variant | Patient ID | Gene  location | Guide RNA sequence | ssODN (sequence) | length |
| --- | --- | --- | --- | --- | --- |
| c.277-560 T>G | P3 | *COL4A5*  Intron 4 | AGGTCAAAGGGCAAATGCAT | A*G*A*TTCTTAGAAGTGAAATGCGAGGTCAAAGGGCAA  ATGCATGGGTAATTTTGCTAG*A*T*A | 60nt |
| c.609-879 A>G | P14 | *COL4A5*  Intron 10 | GCTTTCCTGTAACTTTTTGG | T*G*A*TTTCTGCTTTCCTGTAACTTTTTGGGGGAAGGTG  AGAAAACCCTTTTGTTTGTT*T*T*A | 60nt |
| COL4A5-v5 tag | - | *COL4A5*  Exon 2 | GCTTGCTATGGGTGTTCTCC | T*A*T*AGG CTT GCT ATG GGT GTG GTA AGC CTA TCC CTA ACC CTC TCC TCG GTC TCG ATT CTA CGT CTC CAG GAT CAA AGT G*T*G*A | 82nt |

*Phosphorothioate modifications

| No | Reagent type | Designation | Provider | Identifier | Additional comments |
| --- | --- | --- | --- | --- | --- |
| 1 | **Antibody** | Anti-laminin S/laminin β2 chain, clone CL2979 (mouse monoclonal) | Novus Biologicals | Cat# NBP-42387 | IF (1:100) |
| 2 | **Antibody** | Anti-human collagen IV α3 chain NC1 domain, clone H31 (rat monoclonal) | Chondrex | Cat# 7076 | IF (1:100) |
| 3 | **Antibody** | Anti-human collagen IV α4 chain NC1 domain, clone H43 (rat monoclonal) | Chondrex | Cat# 7073 | IF (1:100) |
| 4 | **Antibody** | Anti-human collagen IV α6 chain NC1 domain, clone H63 (rat monoclonal) | Chondrex | Cat# 7074 | IF (1:100) |
| 5 | **Antibody** | Anti-human collagen IV α5 chain, clone H53 and NC1, clone H52 | Chondrex | Cat# 7078  Cat# 7077 | IF (1:100) |
| 6 | **Antibody** | V5-Tag antibody \| SV5-Pk1, mouse | Bio-rad | Cat# MCA1360 | IF (1:100) |
| 7 | **Antibody** | Rabbit Anti-Claudin-1 (CLDN1), Rabbit Polyclonal Antibody | Bicell | Cat# 00201 | IF (1:100) |
| 8 | **Antibody** | Human GATA-3 Antibody, Goat | R&D systems | Cat# AF2605 | IF (1:100) |
| 9 | **Antibody** | Lotus Tetragonolobus Lectin (LTL), Biotinylated (B-1325-2) | Vector | Cat# B-1325-2 | IF (1:50) |
| 10 | **Antibody** | Anti-laminin alpha 5/LAMA5, Mouse | Abcam | Ab210957 | IF (1:100) |
| 11 | **Antibody** | Anti-laminin beta 1/LAMB1, Mouse | Merk | MAB1921P | IF (1/200) |
| 12 | **Antibody** | E-Cadherin (ECCD), Rat | Life technologies | Cat# B-1900 | IF (1/100) |
| 13 | **Antibody** | Antibody anti-WT1, Mouse (F-6) | Santa Cruz | Cat# sc-7385 | IF (1/100) |
| 14 | **Antibody** | Anti-human collagen IV, Rabbit | Rockland | Cat# 600-401-106-0.1 | IF (1/200) |
| 15 | **Antibody** | Anti-human Nephrin, Sheep | R&D systems | Cat# AF4269 | IF (1/100) |
| 16 | **Antibody** | Anti-human podocalyxin, Goat | R&D systems | Cat# AF1658 | IF (1/500) |
| 17 | **Antibody** | Anti-human ZO-1/TJP1, Rabbit | Invitrogen | Cat# 40-2200 | IF (1/100) |
| 18 | **Antibody** | Anti-human COL4A1-COL4A2 | Developmental Studies Hybridoma Bank  (DSHB) | Cat# M3F7-s | IF (1/100) |

**Table S9. List of the antibodies used in this study**

| Case | Primer | Sequence (5'-3') | Modification | length | Tm | GC | Transcript type | Product size |
| --- | --- | --- | --- | --- | --- | --- | --- | --- |
| P3 | *Forward primer* | CGGGGACAAAAGGGTGATGA | FAM | 20 | 59.96 | 55 | **Wild type** | 177 bp |
|  | *Reverse primer* | TCCACGTTCTCCCTTGGTTC | - | 20 | 59.6 | 55 | **Mutant** | 235 bp |
| P14 | *Forward primer* | CCTGGTCCCACTGGTATACC | FAM | 20 | 58.87 | 60 | **Wild type** | 123 bp |
|  | *Reverse primer* | GGGTCCCTGGAAATTTAAGCC | - | 21 | 58.89 | 52 | **Mutant** | 251 bp |
| P10 | *Forward primer* | GAAAGGAGATCAAGGACCACC | FAM | 21 | 58.01 | 52.38 | **Wild type** | 182 bp |
|  | *Reverse primer* | GAGGACCTGGAGGACCAATAA | - | 21 | 58.52 | 52.38 | **Mutant** | 248 bp |
| P2 & P9 | *Forward primer* | CAGACGATCCAGATTCCCCA | FAM | 20 | 58.88 | 55 | **Wild type** | 286 bp |
|  | *Reverse primer* | GTGTCCTCAAGTCTCCTGCT | - | 20 | 59.03 | 55 | **Mutant** | 103 bp |
| P16 | *Forward primer* | CACCAGGACCAAAAGGAATCA | FAM | 21 | 58.12 | 47.62 | **Wild type** | 170 bp |
|  | *Reverse primer* | ACCAGGAAAACCGGGACTG | - | 19 | 59.55 | 57.89 | **Mutant** | 212 bp |
| P15 & P20 | *Forward primer* | TGGTATCAAAGGTTCTGTGGGA | FAM | 22 | 59.02 | 45.45 | **Wild type** | 187 bp |
|  | *Reverse primer* | ATGACCTCCACCACCTACAG | - | 20 | 58.43 | 55 | **Mutant** | 225 bp |
| P19 | *Forward primer* | GAAGGACCTAAAGGCAACCCT | FAM | 21 | 59.65 | 52.38 | **Wild type** | 284bp |
|  | *Reverse primer* | CCTTTCATGCCTGGGAATCC | - | 20 | 58.59 | 55 | **Mutant** | 330bp |
| *COL4A1* | *Forward primer* | ACTACTCGTACTGGCTGTCC | - | 20 | 58.54 | 55 | **Product size** | 151bp |
|  | *Reverse primer* | GGGATCTGAATGGTCTGGCT | - | 20 | 59.16 | 55 |  |  |
| *COL4A2* | *Forward primer* | CCAGGACAGAAAGGAGACCA | - | 20 | 58.65 | 55 | **Product size** | 112bp |
|  | *Reverse primer* | CAGGTGCCCCAGAGATGTT | - | 19 | 59.31 | 57.89 |  |  |
| *COL4A3* | *Forward primer* | GTCAACACCAGCTCTGATGC | - | 20 | 59.2 | 55 | **Product size** | 167bp |
|  | *Reverse primer* | TCCAGAGAGAAATCCAGCCG | - | 20 | 59.17 | 55 |  |  |
| *COL4A4* | *Forward primer* | CAGGCATGAGAGGACCAGAA | - | 20 | 59.09 | 55 | **Product size** | 162bp |
|  | *Reverse primer* | CGTCTTCTCCTGTGTCACCT | - | 20 | 59.04 | 55 |  |  |
| *COL4A5* | *Forward primer* | CAGACGATCCAGATTCCCCATTGT | - | 24 | 62.50 | 50 | **Product size** | 122bp |
|  | *Reverse primer* | GAACCAGGGGAGGCTAGGGCTTGA | - | 24 | 62.50 | 62 |  |  |
| *HPRT1* | *Forward primer* | TCTTTGCTGACCTGCTGGATT | - | 21 | 59.93 | 47.62 | **Product size** | 151bp |
|  | *Reverse primer* | GTTGAGAGATCATCTCCACCAATTACT | - | 27 | 60.63 | 40.74 |  |  |

**Table S10. Primers used for RT-PCR and RT-qPCR**
